# The parthenogenesis mechanism and venom complement of the parasitoid wasp *Microctonus hyperodae*, a declining biocontrol agent

**DOI:** 10.1101/2022.09.11.507509

**Authors:** Sarah N. Inwood, Thomas W.R. Harrop, Peter K. Dearden

## Abstract

A biocontrol system in New Zealand using the endoparasitoid *Microctonus hyperodae* is failing, despite once being one of the most successful examples of classical biocontrol worldwide. In this study, RNA-seq was used to characterise two key traits of *M. hyperodae* in this system, the venom complement, critical for the initial success of biocontrol, and the asexual reproduction, which influenced the decline. Full characterisation of *M. hyperodae* venom revealed 82 candidate venom transcripts with both signal peptides and significantly higher expression in venom. Among these were many involved in manipulating the host environment to source nutrition for the parasitoid egg, preventing a host immune response against the egg, as well as two components that may stimulate the host’s innate immune system. Notably lacking from this list was calreticulin, as it also had high expression in the ovaries. *In-situ* hybridisation revealed expression was localised to the follicle cells, which may result in the deposition of calreticulin into the egg exochorion. Investigating the asexual reproduction of *M. hyperodae* revealed core meiosis-specific genes had conserved expression patterns with the highest expression in the ovaries, suggesting *M. hyperodae* parthenogenesis involves meiosis and the potential for sexual reproduction may have been retained. Upregulation of genes involved in endoreduplication provides a potential mechanism for the restoration of diploidy in eggs after meiosis.

## Introduction

*Microctonus hyperodae* Loan (Hymenoptera: Braconidae) is an endoparasitoid wasp of significant economic importance in New Zealand (NZ) due to its use as a classical biocontrol agent. *M. hyperodae* was released in the 1990s to combat the Argentine stem weevil (ASW; *Listronotus bonariensis* Kuschel) (Coleoptera: Curculionidae), which caused NZ$200M in annual losses before the establishment of an effective control measure (Prestidge et al., 1991). The wasp stings adult weevils and oviposits into the body cavity, with the host surviving until the wasp larva emerges. While initial parasitism rates were as high as 90% three years post-release (Barker & Addison, 2006), continued monitoring has revealed significant declines in parasitism (Goldson & Tomasetto, 2016; Popay et al., 2011; Tomasetto et al., 2017).

This biocontrol decline is hypothesised to be a result of the ASW evolution of resistance to *M. hyperodae*, due to the initial parasitism rates imposing a strong selective pressure. This would be one of the first examples of evolved resistance to previously successful biocontrol, though genomic investigations of ASW populations are yet to reveal a genetic mechanism for this decline (Harrop et al., 2020). Despite its value as a biocontrol agent in NZ, limited genetic analyses have been performed on *M. hyperodae* thus far, and little is known about traits that play key roles in either the initial success or decline of this system. In this study, we aimed to characterise two such traits in *M. hyperodae*, the venom complement and the asexual reproductive mode.

Parasitoid venoms play key roles in ensuring successful parasitism, and the venom complement of *M. hyperodae* likely contributed to the high parasitism rates initially observed post-release. Parasitoid venom is required to manipulate the host during parasitism, e.g. by preventing immune responses to the parasitoid egg (which is one of the most well-characterised parasitism resistance mechanisms) and providing nutrition for the developing egg (Asgari & Rivers, 2011). Venom is particularly important for koinobiont adult endoparasitoids such as *M. hyperodae*, where the egg is oviposited into the host hemocoel and develops within the surviving adult host, as these manipulations must be long-lasting (Moreau & Asgari, 2015). Investigation of *M. hyperodae* venom has been limited, with eight components identified with dideoxy-sequencing (three of which were subject to mass spectrometry), and their expression patterns across other tissues not characterised (Crawford et al., 2008). An SDS-PAGE gel of *M. hyperodae* venom also contained far more than eight bands (Crawford et al., 2008), with parasitoid wasp venoms often containing 70 or more venom proteins (Danneels et al., 2010; Scieuzo et al., 2021; Sim & Wheeler, 2016; Yan et al., 2016), so this is unlikely to represent the full venom complement.

Unlike venom, the reproduction mechanism of *M. hyperodae* has been detrimental to biocontrol success. While ASW reproduces sexually and has a considerable genetic diversity (Harrop et al., 2020), *M. hyperodae* reproduce asexually via thelytokous parthenogenesis and went through a severe bottleneck upon introduction to NZ (Goldson et al., 1993). Modelling shows this asymmetry in reproduction strategies and genetic diversity contributed to the biocontrol decline and evolution of resistance (Casanovas et al., 2018). However, during the importation and rearing of *M. hyperodae* for release as a biocontrol agent four impotent males were collected (Goldson et al., 1990), indicating *M. hyperodae* has the potential to reproduce sexually in their home range. Sexually reproducing *M. hyperodae* could be used to increase genetic diversity and for selective breeding programs to increase biocontrol efficacy.

To provide a more comprehensive investigation of both *M. hyperodae* venom and parthenogenesis we used RNA-seq to *de novo* assemble a transcriptome and survey gene expression across multiple tissues. A meta-transcriptomic analysis detected viral genes expressed in *M. hyperodae* but did not reveal common parthenogenesis-causing bacteria. Differential gene expression analysis identified both new venom components and improved assembly and annotation of previously characterised components. Further investigation of the expression pattern of calreticulin also revealed this core parasitoid venom component is not venom-specific in *M. hyperodae*, with expression in the follicle cells of the ovaries. Core meiosis genes were present in the transcriptome, with the highest expression in the ovaries indicating *M. hyperodae* asexual reproduction involves meiosis, retaining the potential for sexual reproduction.

## Methods

### Library preparation for RNA-seq

*M. hyperodae* were collected from Lincoln, New Zealand in batches from November to March 2019/2020 and 2020/2021 and were dissected in PBS under a Leica dissection microscope using fine-tipped forceps. Three replicates of twenty pooled individuals were collected for the head, thorax, ovaries, venom gland and reservoir, and remaining abdomen tissue. Three replicates of ten pooled pupae were collected, all three days after cocoon formation. Samples were stored at −80°C, before extraction, and RNA was extracted using a hybrid of Trizol (Ambion) and RNeasy mini kit (Qiagen) methods. RNA purity was assessed using a Nanodrop 2000, and RNA integrity was measured using the Agilent 5200 Fragment Analyzer System, with all samples passing these quality assessments. Samples were then prepared for sequencing using the low-input paired-end (2 × 100bp) Illumina stranded mRNA platform and sequenced on an Illumina HiSeq 2500 by Otago Genomics Facility (https://www.otago.ac.nz/genomics/index.html). An additional head, abdomen and two ovary RNA-seq samples from two previous RNA-seq runs prepared using the same extraction and library preparation protocols were also used in the analyses below.

### Pre-processing of RNA-seq samples for transcriptome assembly

BBDuk v38.90 (Bushnell, 2014) was used to quality trim and remove Illumina sequencing adapters, using default settings with trimq set to 15, and ftl=1 to remove the T overhang added by the Illumina stranded mRNA library platform. FastQC v11.9 was then used to ensure quality trimming was successful and no further issues remained with samples. Kraken2 v2.1.2 (Wood et al., 2019) was then used to taxonomically classify reads against the Kraken standard database (downloaded 17^th^ May 2021), and to identify potentially contaminated samples, given the low input RNA concentrations.

### De novo *M. hyperodae* transcriptome assembly and quality assessment

Before assembly, BBMerge v38.90 (Bushnell et al., 2017) was used to merge overlapping reads using the ‘very strict’ setting. *De novo* transcriptome assembly was performed with Trinity v2.12.0 (Grabherr et al., 2011) using default settings with output from BBMerge for all samples retained after Kraken2 analysis (excluding one contaminated venom replicate). Transcript redundancy was reduced by retaining the longest transcript assembled for each gene. BUSCO v5.2.1 (Simão et al., 2015) was then used to assess transcriptome completeness using the hymenoptera_odb10 lineage. The scripts used for this pipeline are available in an open-source repository at https://github.com/sarahinwood/mh-transcriptome with a Snakemake-managed workflow (Köster & Rahmann, 2012).

### *M. hyperodae* transcriptome annotation

Functional annotation of the *M. hyperodae* transcriptome was performed using the Trinotate pipeline v3.2.0 (Bryant et al., 2017). BlastX v2.9.0 (Altschul et al., 1990) was used against the UniProtKB/SwissProt database (Boutet et al., 2007) downloaded on February 3^rd^ 2020. This was used to generate KEGG (Kanehisa et al., 2012), GO term (Ashburner et al., 2000) and eggNOG (Powell et al., 2012) annotations. Transdecoder v4.0.0 was then used to predict protein-coding regions within transcripts. Transdecoder output was then used for a BlastP v2.9.0 (Altschul et al., 1990) search against the same UniProt database to retrieve the same information as BlastX, by Hmmer v3.1b2 (Finn et al., 2011) to identify protein domains using the Pfam database (Finn et al., 2014) downloaded 3^rd^ February 2020, and by SignalP v4.1 (Petersen et al., 2011) to predict signal peptides. Annotations were loaded into an SQLite database. A Python 3 wrapper for this annotation is available in an open-source repository at https://github.com/sarahinwood/trinotate_pipeline.

### Reciprocal BlastX for viral transcripts

As many insect viruses are not present in the UniProtKB/Swiss-prot database used for Trinotate annotation, an alternate method was used to investigate the virome of *M. hyperodae*. This involved performing a BlastX v2.12.0 search (with an E value of 1E-05 to minimise false-positive results) of all transcripts against the non-redundant (nr) database (Pruitt et al., 2007) downloaded on May 16^th^, 2021. This search was restricted to viral entries in the database by using TaxonKit 0.8.0 (Shen & Ren, 2021) to produce a list of all viral taxonomy identifiers at a species level or below, and restricting the BlastX search to this list using the -taxidlist option. Any gene with a significant BlastX result to a virus was then used in a subsequent BlastX search against the whole nr database to remove genes with better non-viral hits. All transcripts with significant hits that had sequence identity over 90% were then subject to a BlastN v2.12.0 search against the nucleotide database (downloaded March 16^th^, 2022), with an E-value threshold of 1E-05.

### RNA-seq analysis pipeline

All samples were *quasi*-mapped against the length-filtered *M. hyperodae* transcriptome using Salmon v1.5.1 (Patro et al., 2017) with default settings. DESeq2 v1.30.1 (Love et al., 2014) was used to create the DESeqDataSet (DDS) object, by importing Salmon output files using tximport v1.18.0 (Soneson et al., 2016) in R v4.0.4 (R Development Core Team, 2020). A blind variance stabilising transformation (VST) was performed on the DDS object, which was used for principal component analysis (PCA). Differentially expressed genes (DEGs) were identified by filtering DESeq2 results with the arbitrary alpha threshold value of 0.05 for all analyses, and a log fold change magnitude of greater than 1 for Wald tests.

DESeq2 v1.30.1 (Love et al., 2014) was used to investigate differential expression between adult tissue samples. Iterative pairwise Wald tests were carried out comparing the tissue of interest to each other adult tissue separately, using the design ∼Flowcell+Batch+Tissue. This design controlled for the effect of different sequencing runs, and variance between sample batches for each sequencing run, as batches were collected consecutively across two summers and this was shown to cause tighter clustering of some head, thorax, and abdomen samples in the PCA than tissue did (particularly for batch two where the head and thorax are directly on top of one another, supplementary figure 1). For each tissue, results from each comparison were overlapped and genes detected as significantly differentially expressed in all comparisons were retained as a list of tissue-specific genes. Result tables (including log_2_ fold-changes and adjusted P-values) were retained from the venom vs ovary comparisons, after being subset to only contain genes differentially expressed in all pairwise comparisons.

Heatmaps of expression were generated using VST normalised data with pheatmap v1.0.12 (Kolde, 2019) for comparisons of genes of interest, with unsupervised clustering of genes and samples based on expression patterns. Enrichment of gene ontology terms in tissue-specific gene lists was investigated using ClusterProfiler v3.18.1 (Yu et al., 2012).

### Venom-specific analyses

A BlastN v2.9.0 (Altschul et al., 1990) search was carried out for the sequences from Crawford et al. (2008) against the *M. hyperodae* transcriptome database (with an E value of 1E-5), retaining the Trinity ‘genes’ with the best hit to each previously characterised venom component.

BlastX v2.12.0 (Altschul et al., 1990) homology searches of any DEGs without Trinotate annotation, as well as Crawford venom gene transcripts, and all venom DEGs with signal peptides, were performed (with an E value of 1E-5 to minimise false-positive results) against the non-redundant database (Pruitt et al., 2007), downloaded May 16^th^ 2021. Results were first filtered to remove uncharacterised or hypothetical annotations that provided no information on gene function before the result with the lowest E-value (and highest bit score in the case of a tie) was selected for any genes with results.

### Ovary-specific analyses

To identify core meiosis genes, Trinotate BlastX, BlastP and Pfam annotations were searched for the following genes: CORT (cell cycle regulation), REC8 (sister chromatid cohesion), SPO11, MND1, HOP2 and DMC1 (meiotic interhomolog recombination), and MSH4 and MSH5 (crossover resolution) (Schurko et al., 2010; Schurko & Logsdon, 2008; Tvedte et al., 2017). The resulting hits were then filtered based on whether Blast hits or Pfam domains indicated a hit to a meiosis-specific gene or a homolog. Any genes with these meiosis-specific annotations were subject to a BlastX v2.12.0 search against the nr database (with an E value of 1E-5), with expression patterns also investigated. Given a lack of hits to REC8 and DMC1, the *M. hyperodae* transcriptome was the subject to a BlastX v2.12.0 search (with an E value of 1E-5) against sequences for these genes used in a previous investigation of Hymenopteran meiosis genes (Tvedte et al., 2017) to further investigate their presence. Any transcripts with a significant hit were then searched against the non-redundant database with BlastX v2.12.0 (with an E value of 1E-5), and results were filtered to retain the best hits after the removal of hypothetical or uncharacterised hits.

After the presence of these genes in the ovary-specific DEG list was investigated, a likelihood ratio test (LRT) was performed, with the design ∼Flowcell+Batch+Tissue and reduced model of ∼Flowcell+Batch, to determine whether expression patterns of meiosis-specific genes were significantly influenced by tissue. The scripts used for this differential expression analysis pipeline are available in an open-source repository at https://github.com/sarahinwood/mh-rnaseq with a Snakemake-managed workflow.

### Hybridisation chain reaction for Calreticulin expression

Hybridisation chain reaction (HCR) (Choi et al., 2016, 2018) was used to investigate the expression of calreticulin in the ovaries of ten *M. hyperodae*. Dissected ovaries were fixed for five minutes by rocking at room temperature in a 1:1 mix of heptane and 4% formaldehyde in PBS (phosphate buffered saline). The lower heptane layer was replaced with 100% ice-cold methanol and incubated for five minutes. The upper formaldehyde layer was then removed, and tissue was washed in ice-cold methanol three times for five minutes, before being stored at −20 °C.

Hybridisation was performed using HCR version 3 (Choi et al., 2016, 2018). Ovaries were rehydrated in successive five-minute 75%, 50% and 25% methanol washes. Ovaries were then washed 3x for five minutes in PTw (PBS + 0.1% Tween 20). Ovaries were pre-hybridised in 500 µL of 30% probe hybridisation buffer (30% formamide, 5x sodium chloride sodium citrate (SSC)) for 30 minutes at 37 °C. The probe solution, containing 1 µL of calreticulin probe in 500 µL probe hybridisation buffer, was then added to tissues and incubated overnight at 37 °C. Excess probes were removed by washing with probe wash buffer (30% formamide, 5X SSC, 9 mM citric acid (pH 6.0), 0.1% Tween, 50 μg/mL heparin) four times for 15 minutes each at 37 °C. Samples were then washed with 5X SSCT (5X SSC, 0.1% Tween 20) three times for 5 minutes each at room temperature.

In this study, Calreticulin was used with fluorophore 488. The hairpin mixture was prepared by incubating 2 µL of each 3M hairpin stock at 95 °C for 30 seconds in separate tubes, then left to cool at room temperature for 30 minutes. To prevent bleaching hairpins, all subsequent steps were carried out in the dark. 100 µL of amplification buffer (5X SSC, 0.1% Tween 20, 10% dextran sulphate) was added to hairpins, and the amplification buffer with tissue was replaced with the hairpin mixture and incubated overnight at room temperature. The hairpin solution was then removed, and tissues were washed with 5x SSCT as follows; 2x 5 minutes, 2x 30 minutes, 1x 5 minutes, and then stored in 70% glycerol. Tissues were mounted on slides and imaged using an Olympus BX61 Fluoview FV100 confocal microscope with FV10-ASW 3.0 imaging software.

## Results

### *M. hyperodae* transcriptome assembly

Before assembly, taxonomic classification of reads was performed with Kraken 2 to ensure samples were not contaminated during preparation. We expected samples would contain predominantly unclassified reads as the Kraken2 database does not contain insects. Kraken 2 revealed most samples had between 90.0% to 96.8% of reads unclassified, though detected significant contamination in one venom replicate with 29.8% reads unclassified, 52.1% reads classified as bacterial, and 15.0% reads classified as human. This venom replicate (Mh_venom_3) was removed from all subsequent analyses.

Sequencing from the remaining samples generated 36.8-68.7 million reads per sample, with trimming retaining over 99.7% of reads (Supplementary table 1). *De novo* assembly with Trinity resulted in 202,655 transcripts sorted into 120,072 ‘genes’ with a GC content of 32.9%. BUSCO analysis indicates that our transcriptome has a high level of completeness though many BUSCO genes were duplicated (C:94.0% [S:5.5%, D 88.5%] F: 2.9%, M:3.1%). This was reduced when the assembly was filtered to retain only the longest isoform per gene without a large decrease in complete BUSCO genes (C:92.2% [S:81.6%, D:10.6%), F:3.6%, M:4.2%) indicating BUSCO redundancies were due to the assembly of multiple transcript isoforms per gene. Therefore, further analyses used the longest isoform per gene only.

A BlastX search against the UniProtKB/Swiss-prot database as part of the Trinotate pipeline found significant hits for 21.5% of Trinity genes. Transdecoder predicted complete coding sequence in 18.7% of genes, and SignalP predicted signal peptides in 5,430 of these genes. Significant hits to Pfam protein domains were found for 11.6% of genes, with associated GO terms for 7.4%.

### Preliminary investigation of *M. hyperodae* microbiome and virome

Parthenogenesis in insects can be induced by intracellular bacteria, such as those in the *Wolbachia, Cardinium* and *Rickettsia* genera (Ma & Schwander, 2017), with the potential for sexual reproduction retained (e.g. Arakaki et al., 2000; Stouthamer et al., 1990). As well as the impact of the microbiome on sexual reproduction, the virome of parasitoids can play a critical role during parasitism, where endogenous viral genes or proteins from polydnaviruses (PDVs) or virus-like particles (VLPs), and/or exogenous viral infections which can assist in host manipulation (Drezen et al., 2017; Ye et al., 2018). We investigated the bacterial and viral content of reads and the assembled transcriptome. While our libraries were prepared with poly(A) enrichment, such enrichment during RNA library preparation did not prevent the detection of *Wolbachia* infection in *Nasonia vitripennis* ovaries in previous studies (Sim & Wheeler, 2016). Kraken2 classified 4.9% of reads from the remaining samples, with 2.0% bacterial and 0.1% viral. Bacterial reads were predominantly classified as the following phyla: *Proteobacteria* (0.72%), *Firmicutes* (0.48%), *Actinobacteria* (0.25%), *Bacteroidetes* (0.14%), *Cyanobacteria* (0.04%), *Fusobacteria* (0.01%), *Spirochaetes* (0.01%), and *Tenericutes* (0.01%). The *Wolbachia, Rickettsia* and *Cardinium* genera, all known to have caused parthenogenesis in other insects (Ma & Schwander, 2017), have 0.0% reads classified to them (15528, 2445 and 117 reads respectively), and there were no Trinotate annotations from these genera despite their inclusion in the database used for annotation. This is consistent with previous failed attempts to revert asexuality in *M. hyperodae* with antibiotic and heat treatments (Phillips, 1995).

Investigating viral gene content in the *M. hyperodae* transcriptome using a reciprocal BlastX search found 132 genes with significant hits. Many hits were to dsDNA viruses and most had protein sequence identity below 80%, unlikely to represent detection of characterised viruses (Supplementary figure 1, Supplementary table 2). This viral gene search did not reveal any strong PDV gene candidates or other viral genes with expression patterns specific to the venom and/or ovaries that could have indicated a similar role in parasitism. A genome assembly would be required to more extensively survey for endogenous viral elements.

Of the 16 BlastX hits with protein sequence identity above 90%, representing detection of previously identified viruses, nine were from *Iflaviridae*, a viral family known to infect insects. A BlastN search of these transcripts revealed they represent fragmented genomes for Deformed wing virus (DWV) and Moku virus (Supplementary table 3), which have not previously been described as infecting parasitoid wasps. Mean transcripts per million (TPM) were highest for the longest contig of the viral genome, at 2.73 for DWV and 5.53 for Moku virus. Both DWV and Moku virus are known to infect Hymenoptera (Dalmon et al., 2019; Highfield et al., 2020; Mordecai et al., 2016; Wilfert et al., 2016). DWV is well characterised in its negative impacts on honeybees (Koziy et al., 2019), though the impacts of DWV infection in other host species, particularly when it is not vectored by *Varroa* leading to high viral titres, are not known. The impact of Moku virus infection, in general, is also unknown.

### Differential gene expression analysis

Salmon mapping rates against the length-filtered transcriptome ranged between 79.9% and 87.2%. Principal component analysis (PCA) revealed the combination of PC1 and PC2 (which together accounted for 77% of the variance between samples) grouped samples based on tissue identity (Supplementary figure 1). The iterative Wald test results resulted in 782 DEGs in venom (604 with BlastX annotation), and 585 DEGs in the ovaries (412 with BlastX annotation) (Supplementary tables XX). GO enrichment analysis with ClusterProfiler detected significant enrichment of two gene ontology (GO) terms and six Pfam protein domains in the venom list and two GO terms and three Pfam domains in the ovary list (Supplementary figures 3-4).

### Improved characterisation of *M. hyperodae* venom reveals further venom components & expression of calreticulin in the ovaries

As venom components are secreted, they are expected to contain signal peptides in their amino acid sequence. Of the 741 venom DEGs upregulated in venom samples, 83 were predicted to encode a signal peptide by SignalP, 64 of which had BlastX annotations (Supplementary table 4). The absence of a signal peptide on a putative venom gene could be a result of incomplete transcript assembly, therefore the number of venom genes with signal peptides may be higher than this.

A BlastN search was used to identify the best transcriptome hits for the previously sequenced *M. hyperodae* venom components, given their expression patterns had not been investigated. The best transcript hit to venom protein 5 (TRINITY_DN1215_c1_g1_i10) was removed at this stage as Blast results showed it was a chimeric transcript, and the second-best hit instead retained. The transcript hits for venom proteins 3, 4, 5 and 10 were all in the venom-specific protein list, giving more certainty to their function in *M. hyperodae*. Venom protein 1 is still considered a venom-specific candidate as it was only excluded from this list due to an adjusted P-value of 0.08 between venom and ovary samples (Table 1**Error! Reference source not found**.) which may have been lower with an additional venom replicate, and was one of the most abundant proteins in *M. hyperodae* venom (Crawford et al., 2008). Investigation of both these previously identified as well as newly characterised *M. hyperodae* venom-specific gene annotations revealed candidates involved in processes commonly manipulated during parasitism.

**Table 1.**
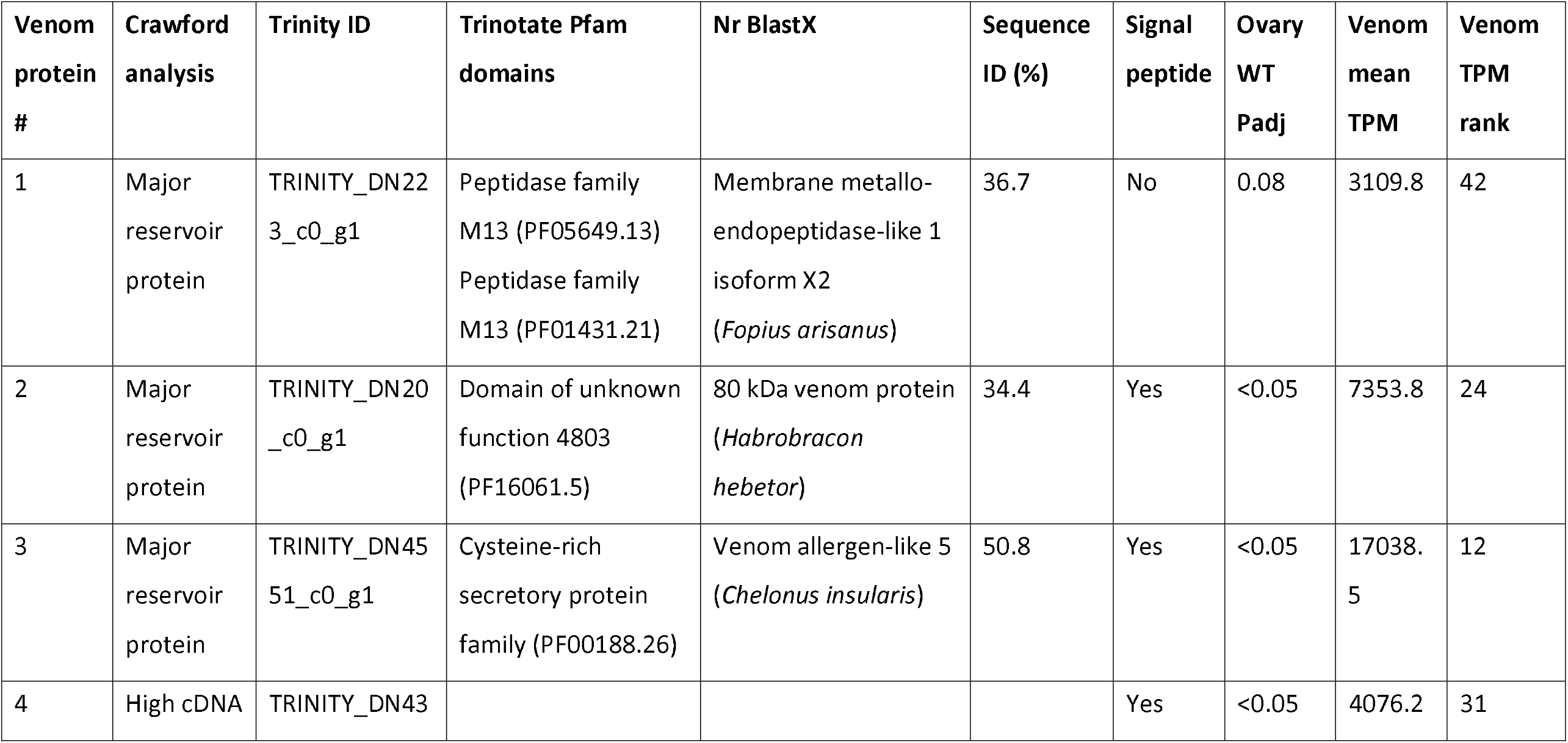
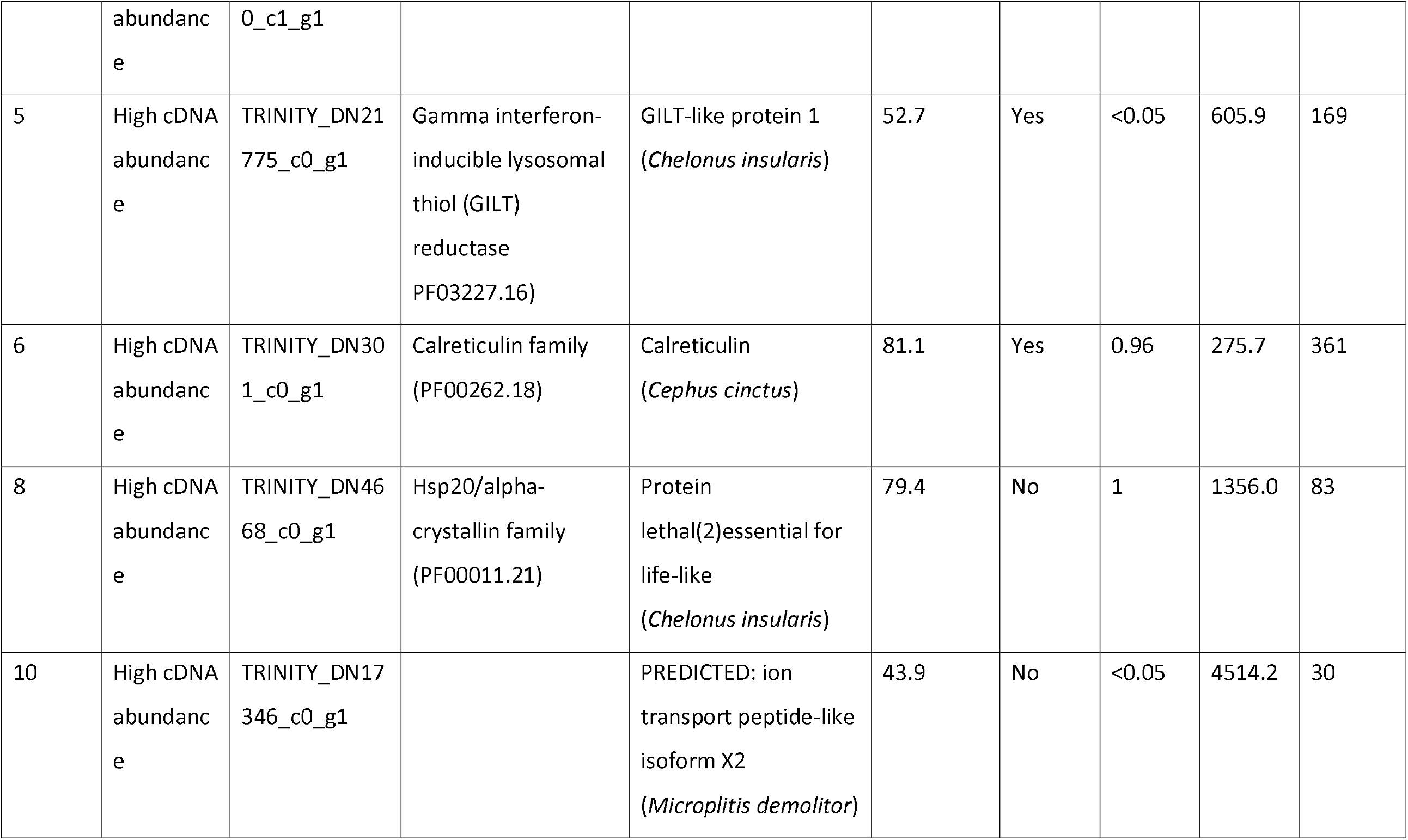
Trinotate Pfam, BlastX and SignalP results for Crawford *M. hyperodae* venom Trinity genes. Venom protein numbers refer to those from Crawford et al. (2008), Trinity IDs are those identified as best hits for Crawford genes using BlastN, Ovary WT Padj the adjusted P value for a differential expression Wald test between venom and ovary samples, mean TPM the mean of TPM calculated by Salmon for the two venom samples and venom TPM rank the placement of each gene in a list ordered from highest to lowest TPM for DEGs upregulated in venom.

| Venom protein # | Crawford analysis | Trinity ID | Trinotate Pfam domains | Nr BlastX | Sequence ID (%) | Signal peptide | Ovary WT Padj | Venom mean TPM | Venom TPM rank |
| --- | --- | --- | --- | --- | --- | --- | --- | --- | --- |
| 1 | Major reservoir protein | TRINITY_DN223_c0_g1 | Peptidase family M13 (PF05649.13)<br>Peptidase family M13 (PF01431.21) | Membrane metallo-endopeptidase-like 1 isoform X2 (*Fopius arisanus*) | 36.7 | No | 0.08 | 3109.8 | 42 |
| 2 | Major reservoir protein | TRINITY_DN20_c0_g1 | Domain of unknown function 4803 (PF16061.5) | 80 kDa venom protein (*Habrobracon hebetor*) | 34.4 | Yes | <0.05 | 7353.8 | 24 |
| 3 | Major reservoir protein | TRINITY_DN4551_c0_g1 | Cysteine-rich secretory protein family (PF00188.26) | Venom allergen-like 5 (*Chelonus insularis*) | 50.8 | Yes | <0.05 | 17038.5 | 12 |
| 4 | High cDNA | TRINITY_DN43 |  |  |  | Yes | <0.05 | 4076.2 | 31 |
|  | abundance | 0_c1_g1 |  |  |  |  |  |  |  |
| 5 | High cDNA abundance | TRINITY_DN21775_c0_g1 | Gamma interferon-inducible lysosomal thiol (GILT) reductase (PF03227.16) | GILT-like protein 1 (*Chelonus insularis*) | 52.7 | Yes | <0.05 | 605.9 | 169 |
| 6 | High cDNA abundance | TRINITY_DN301_c0_g1 | Calreticulin family (PF00262.18) | Calreticulin (*Cephus cinctus*) | 81.1 | Yes | 0.96 | 275.7 | 361 |
| 8 | High cDNA abundance | TRINITY_DN4668_c0_g1 | Hsp20/alpha-crystallin family (PF00011.21) | Protein lethal(2)essential for life-like (*Chelonus insularis*) | 79.4 | No | 1 | 1356.0 | 83 |
| 10 | High cDNA abundance | TRINITY_DN17346_c0_g1 |  | PREDICTED: ion transport peptide-like isoform X2 (*Microplitis demolitor*) | 43.9 | No | <0.05 | 4514.2 | 30 |

### Parasitoid egg nutrition

After parasitism of a host by a koinobiont endoparasitoid wasp, it is critical for the survival of the egg to source nutrients required for development from the host’s tissue, while avoiding killing the host (Moreau & Asgari, 2015). *M. hyperodae* venom candidates for this nutritional sourcing include a venom acid phosphatase, found in many other parasitoid venoms, such as *N. vitripennis* (Danneels et al., 2010), *Leptopilina boulardi* (Colinet et al., 2013) and *Pachycrepoideus vindemmiae* where it was hypothesised to play a role in sourcing nutrients from host hemolymph (Yang et al., 2020). *M. hyperodae* venom also contains several lipases, one of which has a signal peptide and has significant enrichment of the lipase domain (Supplementary figure 3). Lipases are generally important in digestion, processing of dietary lipids and lipid transport (Sahu & Birner-Gruenberger, 2013) and are a common parasitoid venom component. Changes in host lipid metabolism have been observed following parasitism by both *N. vitripennis* (Rivers & Denlinger, 1995) and *Euplectrus* parasitoids (Nakamatsu & Tanaka, 2004), with this manipulation likely to provide a nutritional source for the developing egg.

There are 15 proteases with significantly higher expression in venom, including a serine protease (discussed below), two furin-like proteases, aminopeptidase-N, and angiotensin-converting enzyme-like. Both aminopeptidases and angiotensin-converting enzymes have also been found in the venom of the parasitoid *Torymus sinensis* (Scieuzo et al., 2021) and may assist in manipulating host tissue to provide nutrition for the parasitoid egg and/or degradation of host extracellular matrix to increase venom permeabilization (Vaiyapuri et al., 2010).

### Venom toxins

*M. hyperodae* venom protein 10 is a likely venom toxin, with the best transcript hit annotated as ion transport peptide-like (ITPL) isoform X2, belonging to the Crustacean CHH/MIH/GIH neurohormone family. Ion-transport peptide (ITP) family peptides are neurohormones, well characterized in crustaceans where they play many roles (Webster et al., 2012). ITP peptides have been found in venoms of some spiders, ticks, wasps, and centipedes where they act as toxins called helical arthropod-neuropeptide-derived (HAND) toxins (Undheim et al. 2015; McCowan and Garb 2014). ITPLs have been found in the venom of parasitoid wasp Tetrastichus brontispae where they are also hypothesised to function as toxins (Liu, Xu, et al., 2018), though there is no more specific indication of function beyond this. Alongside this toxin is a peroxiredoxin, which have been found in two parasitoid venoms previously with unclear function (Perkin et al., 2015), but are thought to be involved in structural or functional diversification of toxins in snake venom (Calvete et al., 2009).

### Parasitism of an adult host requiring alternate venom function

As most parasitoids attack the egg or larval stages of their hosts, parasitoid venom components have not been investigated as having roles in affecting an adult host. This may indicate that known venom components include only some of the functions that venoms play during parasitism, or that the functions of venom components have diversified between parasitoids that attack egg/larval stages and those which attack adults. *M. hyperodae* venom contains two chitinases, enzymes that are capable of disrupting and digesting chitin, the major constituent of insect exoskeletons (Kramer & Koga, 1986). Hypotheses of venom function for chitinases to date are specific to parasitoids of egg/larval host stages, where they play a role in disrupting development (Cônsoli et al., 2005). It may instead play a role in breaking physical barriers to better source nutrition for the developing egg or to facilitate the spread of venom and other injected factors.

*M. hyperodae* venom is here also shown to contain metalloproteases (e.g., venom protein 1), a common parasitoid and snake venom component, with significant enrichment of the metalloendopeptidase activity GO term and two metalloprotease domains in the venom-specific gene list (Supplementary figure 3). In parasitoids, metalloproteases have been shown to alter host development (Z. Lin et al., 2019; Price et al., 2009), though with *M. hyperodae* attacking adult ASW such alteration does not occur. Parasitoid metalloproteases may also inhibit haemocyte accumulation, thereby preventing encapsulation of the parasitoid egg (Danneels et al., 2010; Parkinson et al., 2002) providing an alternative function for this venom component.

### Manipulation of the innate immune system

With improved assembly and annotation of the best hit for *M. hyperodae* venom protein 5, the function of this venom component in venom is clearer. The transcript is predicted to contain a Gamma interferon-inducible lysosomal thiol (GILT) reductase domain and had a significant Blast hit to GILT-like protein 1. GILT-like proteins have been found in the venom of the parasitoid *Pteromalus puparum* (Yan et al., 2016). GILT-like 1 in D*rosophila melanogaster* is involved in immune responses to bacterial infection, with RNAi knockdown resulting in increased bacterial loads 24 hours after infection (Kongton et al., 2014).

Alongside this, another venom-specific gene has a hit to waprin-Thr1-like proteins from several parasitoids and other hymenopteran species, with the best hit from *Diachasma alloeum* with 52.6% amino acid sequence identity, though has not been documented in any parasitoid venoms thus far. Snake venoms contain waprin toxins which have been demonstrated to have roles in protease inhibition, the innate immune system and antimicrobial activity (Hagiwara et al., 2003; Nair et al., 2007). A waprin-Thr1-like protein was found in a proteo-transcriptomic analysis of the wood wasp *Sirex nitobei*, where it was *hypothesised* to be involved in preventing bacterial growth during colonization of a symbiotic fungus. Therefore *M. hyperodae* venom contains two different components that act to stimulate the innate immune system, to potentially prevent bacterial infection of the host through the wound created during oviposition.

### Prevention of cellular immune response to parasitoid egg

One of the critical roles of venom components, as well as PDVs in some parasitoids, is to prevent the host immune system from recognising and mounting a cellular immune response against the parasitoid egg. Many *M. hyperodae* venom-specific genes have functions in other parasitoid venoms involved in the manipulation of this immune response. This includes cathepsin, which is a protease involved in the regulation of autophagy in lepidopteran insects (Saikhedkar et al., 2015). Cathepsin has *hypothesised* roles in parasitoid venoms of blocking host immunity, production of venom components and fat body degradation for nutrient mobilisation (Becchimanzi et al., 2017; Heavner et al., 2013).

There are multiple venom components with roles in disrupting the prophenoloxidase (PPO) cascade, which plays a critical role in the melanisation of parasitoid eggs by the host. *M. hyperodae* venom contains a superoxide dismutase (SOD), which has been demonstrated in the venoms of parasitoids L. boulardi and Scleroderma guani to prevent melanization of the parasitoid egg by disrupting the PPO cascade (Colinet et al., 2011; Liu, Huang, et al., 2018). There is also a venom serine protease, and two serine protease inhibitors (serpins) annotated as antichymotrypsin and chymotrypsin inhibitor. Serine proteases have been found in a wide variety of parasitoid venoms and can play a critical role in preventing melanization of the parasitoid egg by blocking prophenoloxidase (PPO) cascade activation (Thomas & Asgari, 2011; Zhang et al., 2004). Serpins again have been found in many venoms, where they form complexes with serine proteases and assist in the regulation of the PPO cascade to prevent melanisation of the parasitoid egg (Gettins, 2002; Kanost & Gorman, 2008; Yan et al., 2017).

### Calreticulin is not venom-specific in *M. hyperodae*

Notably missing from the venom-specific gene list given its presence in many parasitoid venoms, is calreticulin. *M. hyperodae* venom protein 6 was annotated as calreticulin but it was excluded from the venom-specific gene list due to having no significant difference in gene expression between the venom and ovaries (Table 1). Calreticulin is one of the best-studied parasitoid venom components, with RNAi knockdown in *N. vitripennis* revealing it prevents the host immune system from responding to the parasitoid egg (Siebert et al., 2015; Wang et al., 2013; Zhang et al., 2006).

Adjusted P-values for both venom proteins 6 and 8 are above 0.9, with expression of these genes comparable to or higher in ovary samples (Figure 1**Error! Reference source not found.**). This highlights the benefit of performing a comparison to expression levels in other tissues. The expression patterns of venom proteins 6 and 8 indicate they may either have multiple functions in different tissues or be involved in the same function in venom glands, the venom reservoir, and the ovaries. Both genes also appear to have moderate expression in pupae, while most other venom-specific genes have little to none.

**Figure 1.**
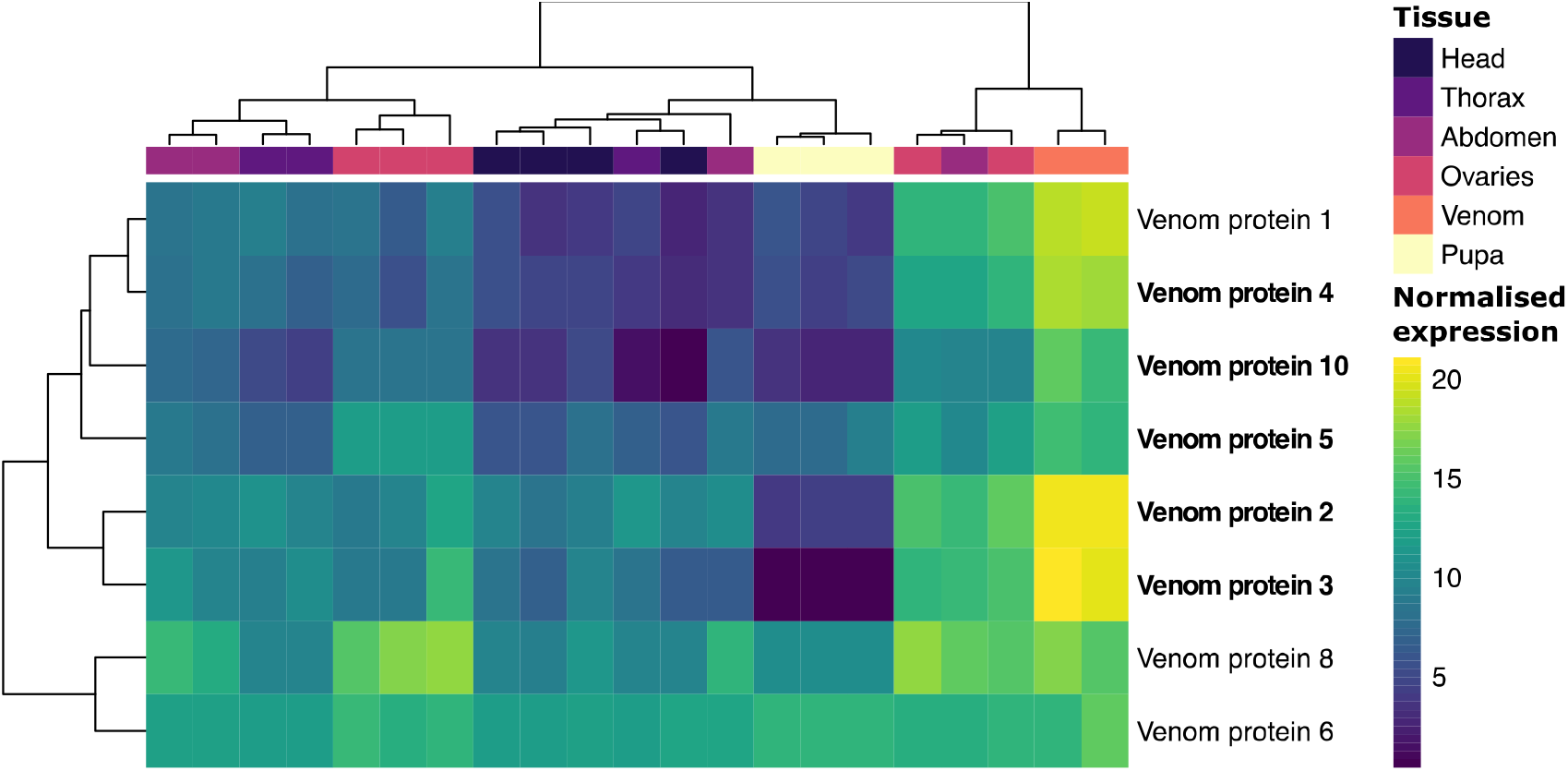
A clustered heatmap showing VST normalized expression for the best hits to previous *M. hyperodae* venom genes from Crawford et al. (2008). Genes significantly differentially expressed in all pairwise Wald tests and the LRT test are bolded. Pupa samples are included in the heatmap, to indicate continued expression of venom proteins 6 and 8 in pupa, though were not included in the differential expression tests.

Given the critical role that calreticulin plays in preventing the host immune response in other parasitoid species, we investigated its RNA expression pattern in the venom gland and ovaries. HCR revealed that calreticulin RNA in the venom gland is in cells on the outside layer arranged around the central lumen (Figure 2), while no expression was detected in the venom reservoir (not shown). This is as expected given a previous histological examination of *M. hyperodae* venom apparatus (Crawford et al., 2006).

**Figure 2.**
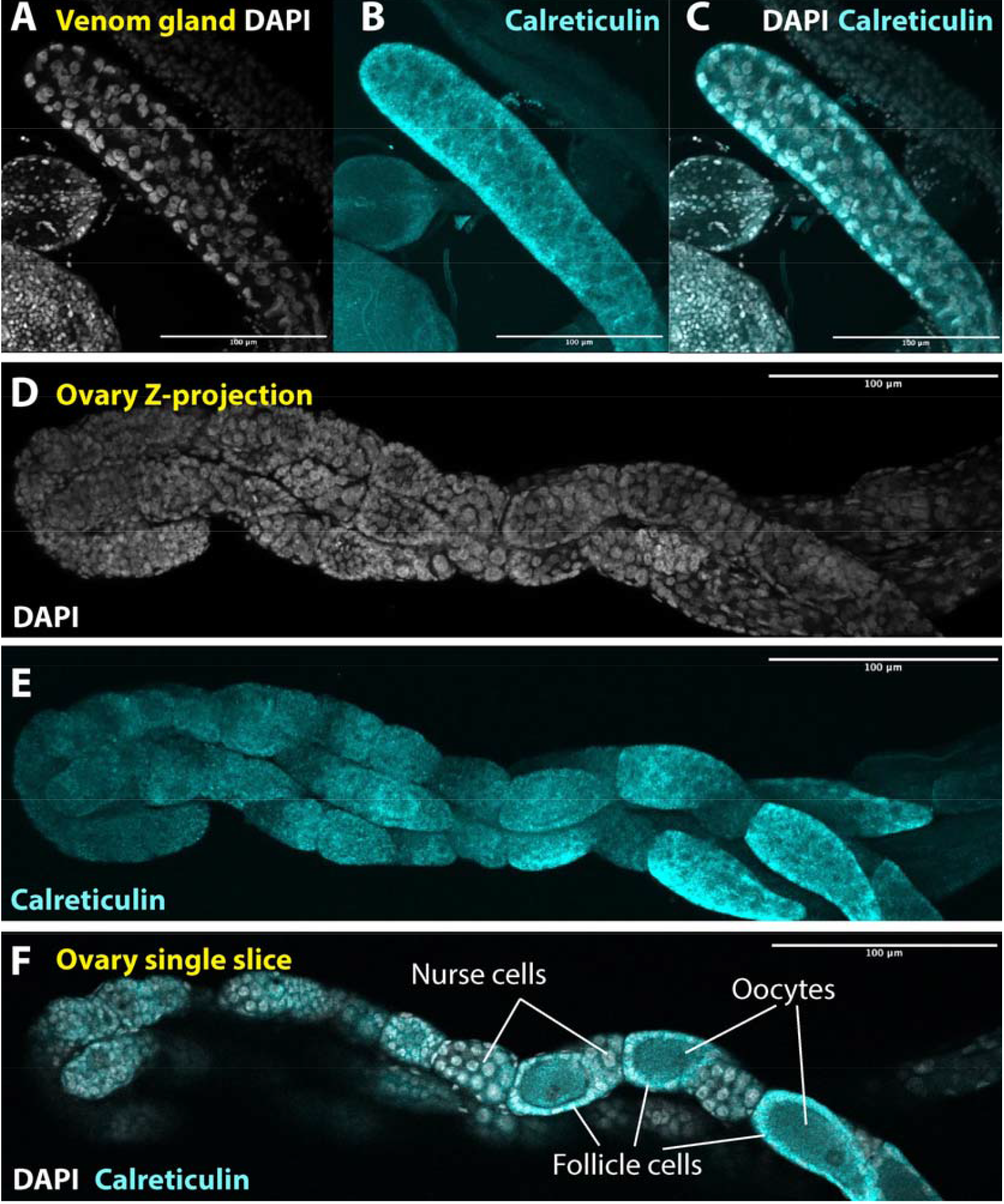
Calreticulin expression in the venom glands and ovaries of *M. hyperodae*, as determined using hybridisation chain reaction.

In the ovaries, calreticulin expression is largely restricted to follicle cells surrounding eggs and was not detected in the oocyte or nurse cells. Calreticulin expression has been detected in the ovaries of N. vitripennis (Sim & Wheeler, 2016), P. puparum (Wang et al., 2013), Cotesia rubecula (Zhang et al., 2006) and Toxoneuron nigriceps (Laurino et al., 2016), though expression patterns across the different cell types in the ovaries have not been investigated before. In these examples it was concluded that such expression may be due to calreticulin being involved in basic cellular processes, for example as a molecular chaperone, or due to its role as a core immune response gene.

Eggs from the parasitoid Hyposoter didymator can evade encapsulation without their associated PDV due to proteins present on the egg exochorion, which are acquired from follicle cells (Dorémus et al., 2013). Proteins with similarity to a venom serpin that prevented melanization of the parasitoid egg have also been detected on the egg surface in Cotesia chilonis, with transcription of this egg serpin mainly in follicle cells (Teng et al., 2021). While calreticulin was not one of the proteins detected in these analyses, this suggests an alternate role for calreticulin expression in the follicle cells of *M. hyperodae* ovaries, where calreticulin protein may be deposited onto the M. *hyperodae* egg exochorion to assist in evading the host immune system.

### Ovary gene expression provides insight into the mechanism of parthenogenesis

The ovary DEG list contained 198 genes with significantly higher expression in the ovaries (Supplementary table 5), many of which were involved with various processes during cell cycle turnover as would be expected during gametogenesis. This included ten genes involved in the Piwi-interacting RNA (piRNA) silencing pathway, including two ovary-specific genes annotated as piwi-like protein SIWI, one as Argonaute-3 (AGO3), three RNA helicases, and one as protein vreteno. piRNA silencing is critical for transposable element silencing in the germline of the sexually reproducing D. melanogaster, with loss of this piRNA silencing causing severe gametogenesis defects (H. Lin & Spradling, 1997).

Ten ovary-specific genes in our dataset are involved directly in cell cycle progression. These include genes involved in the regulation of the G1 to S transition during the cell cycle (CCNE, a cyclin), chromosome condensation (CND2, RCC1, HMGI-C), sister chromatid cohesion (ESCO2, STAG1), chromosome segregation (ESPL1), DNA replication and regulation (DPOA, DPOD), and post-replicative DNA break and mismatch repair (MCM9). This is indicative of gametogenesis occurring in the ovaries, though none of these genes are specific to mitosis or meiosis. The list also included a gene annotated as MARF1, a meiosis regulator and mRNA stability factor, which may indicate meiosis is occurring during *M. hyperodae* gametogenesis.

There are also three DEGs (MCM5, AGO and ESG) involved in endoreduplication, where cells repeat the S phase of mitosis so that DNA is replicated without cell division occurring, increasing cell ploidy. Endoreduplication in *Drosophila* follicle and nurse cells in the ovary is critical for egg production, with disruption leading to sterility (Lilly & Sptadling, 1996; Maines et al., 2004).

### Meiosis-specific genes

To determine whether *M. hyperodae* has retained the potential for sexual reproduction, it is key to determine the mode and cause of parthenogenesis. Parthenogenesis can be broken down into two main mechanisms, automixis and apomixis. Automixis relies on meiosis to produce eggs, using one of several strategies to restore eggs to diploidy, while apomixis involves the production of eggs without meiosis occurring (Tvedte et al., 2019). These two mechanisms can be distinguished by the expression of meiosis-specific genes, which are required for and only expressed during meiosis (Ramesh et al., 2005). In an organism using apomixis, there is no evolutionary constraint acting on genes with meiosis-specific functions, as demonstrated in model organisms, and it is expected that the sequence and function of these genes would not be conserved (Schurko et al., 2010; Schurko & Logsdon, 2008). Loss of these meiosis-specific genes leads to obligative parthenogenesis, where an organism can no longer reproduce sexually, in contrast to facultative parthenogenesis where an organism retains the potential for sexual reproduction.

To identify the mode of parthenogenesis in *M. hyperodae* we identified these meiosis-specific genes in the *M. hyperodae* transcriptome and investigated their expression patterns. Investigation of Trinotate annotations and a subsequent BlastX search for DMC1 and REC8 revealed hits for all genes present in the *M. hyperodae* transcriptome (Table 2). Multiple transcripts were found with Trinotate annotations for MND1 and REC8. Both had one transcript over 2000 bp which was retained for analysis, with other transcript hits below 550 bp and incomplete.

**Table 2.**
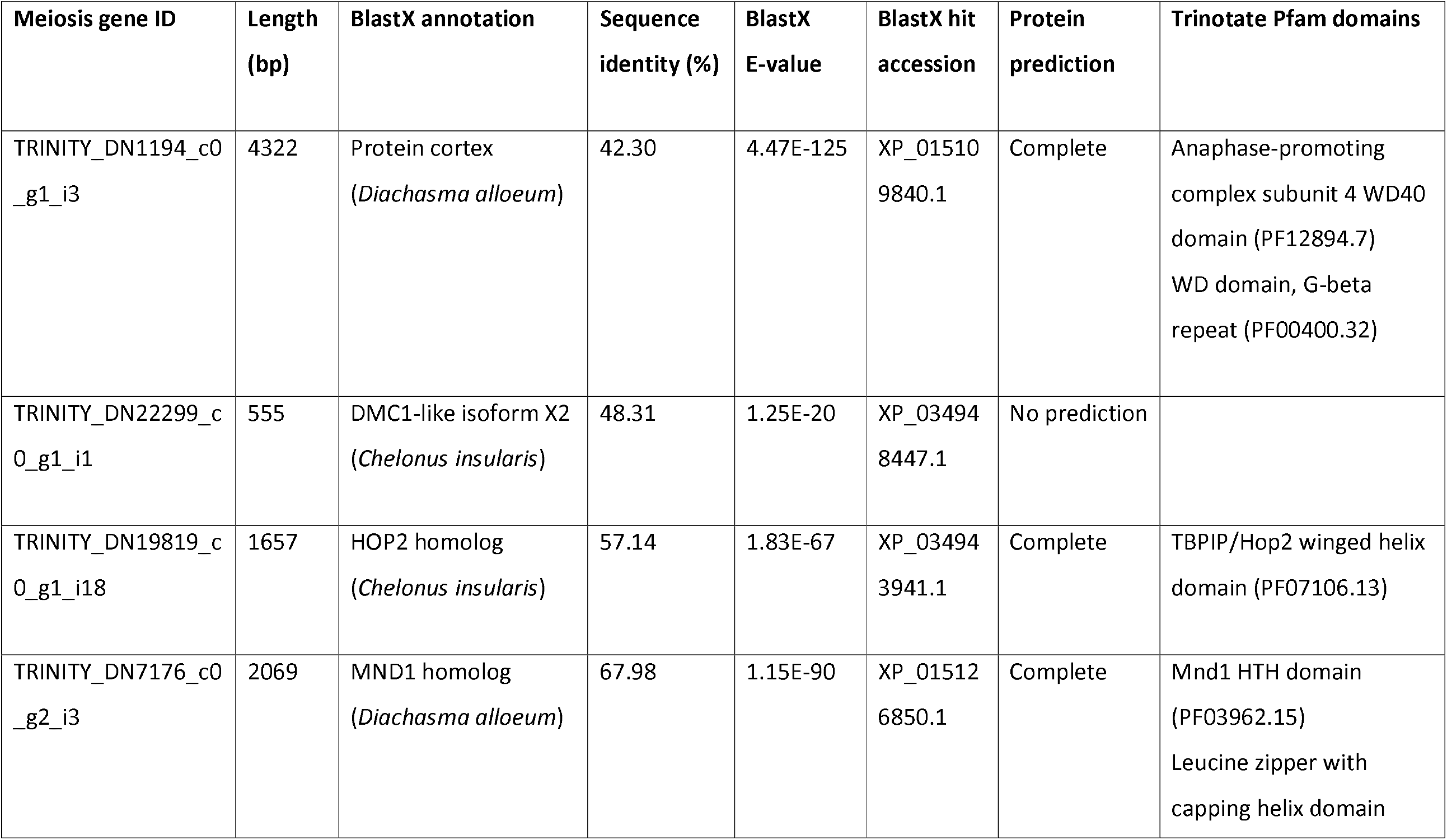
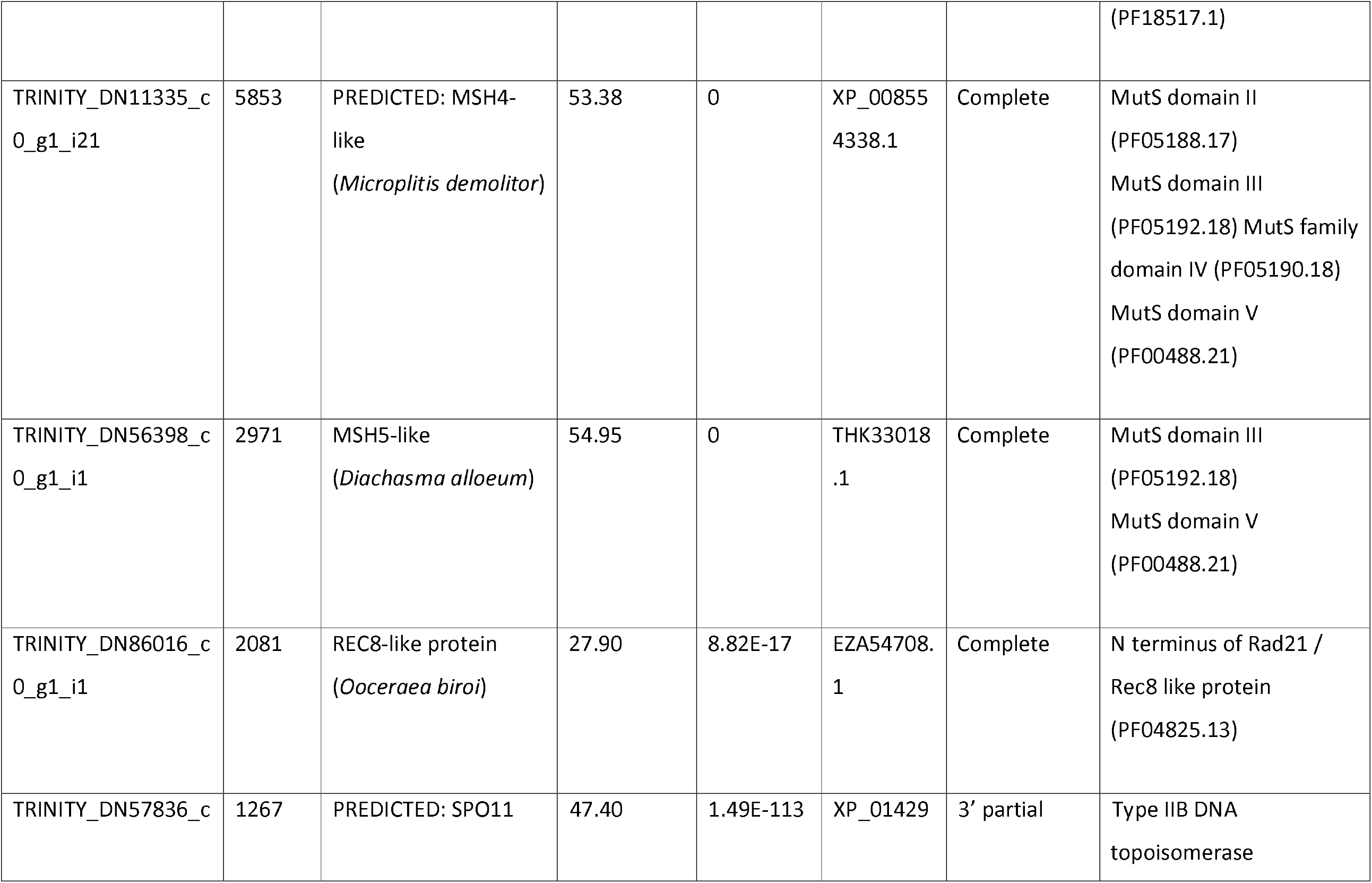
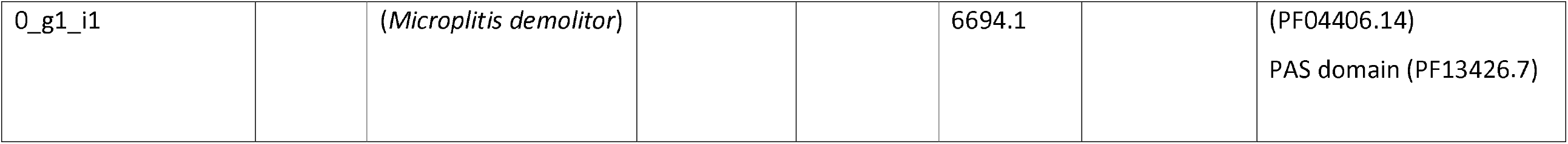
BlastX annotation, Transdecoder protein prediction status, and Pfam results for genes annotated as core meiosis genes.

All but DMC1 had a transcript with a Transdecoder protein prediction and were predicted to contain Pfam domains consistent with the gene, providing further confidence in the identification of these genes. The DMC1 transcript is short and likely incomplete, and an investigation of its expression pattern reveals much lower expression compared to the other meiosis-specific genes (Figure 3). DMC1 has previously been demonstrated to be absent in D. melanogaster and most Hymenoptera investigated, indicating it is not required for meiosis in these species (Tvedte et al., 2017).

**Figure 3.**
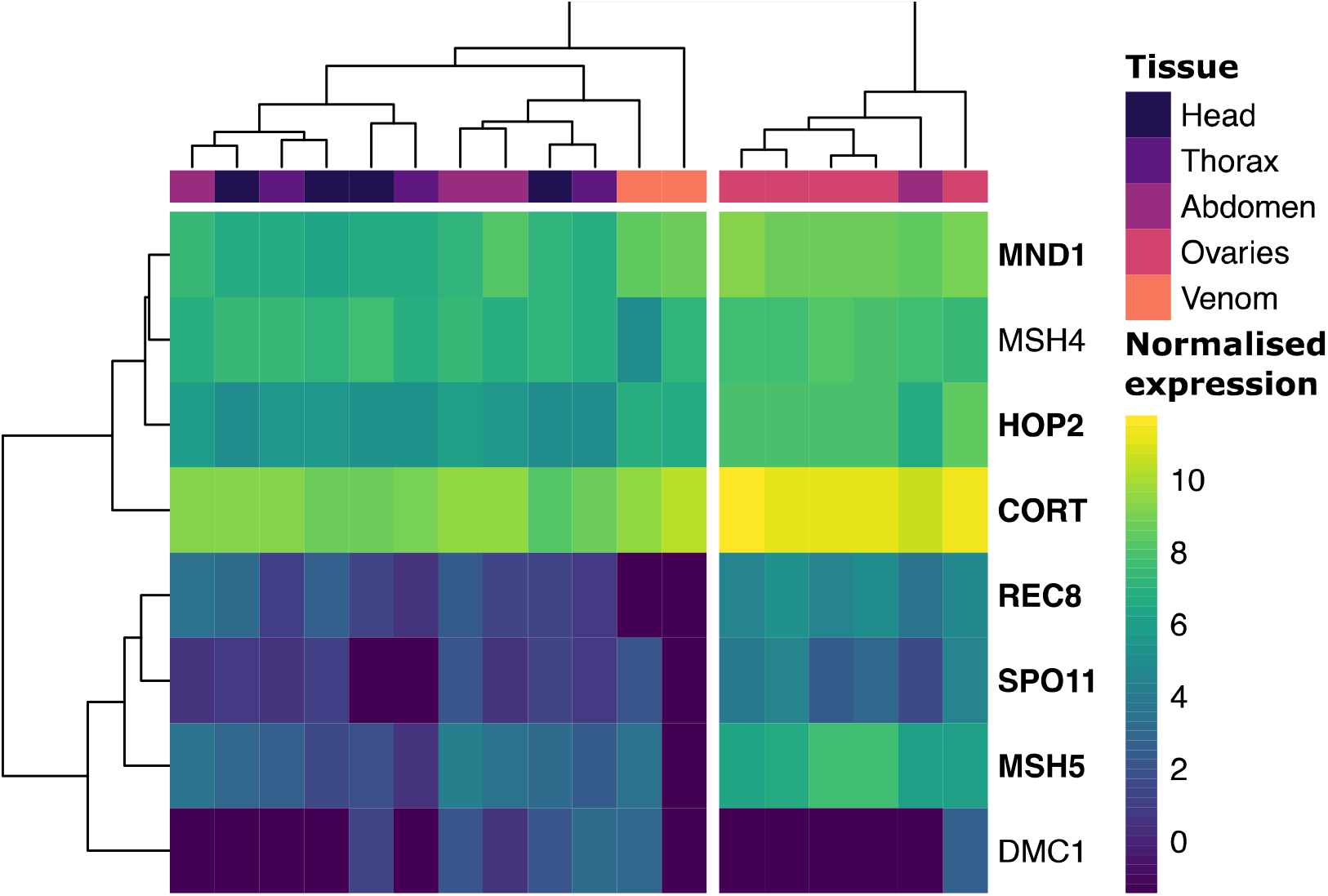
A clustered heatmap showing VST normalized expression for all meiosis-specific genes identified in the *M. hyperodae* transcriptome. Genes that were significantly differentially expressed in the tissue LRT analysis are indicated in bold.

None of the meiosis-specific genes had significant results in all iterative pairwise Wald tests against other tissues, with some having no significant results against any tissue, and others not against venom and/or abdomen samples. However, six of the eight genes were significant in the tissue LRT test (with 28,406 total DEGs from this test) indicating that their expression pattern was significantly influenced by tissue type (Supplementary table 6).

When samples were clustered in an unsupervised manner based on meiotic gene expression, all ovary samples clustered together (with an abdomen sample), against all remaining tissues (Figure 3). The heatmap also shows that the meiosis-specific genes which were significant in the LRT test have the highest expression in the ovaries. The inclusion of an abdomen sample within this clade might be explained by incomplete dissection of ovarian tissue from this sample.

## Discussion

Given the significant economic value of *M. hyperodae* as a biocontrol agent for the ASW in New Zealand, we need a better understanding of key traits that influence the success and later failure of this biocontrol system. Here we have characterised the full venom complement of *M. hyperodae*, and investigated the expression pattern of calreticulin, demonstrating expression in the follicle cells which may serve to protect the egg from the host immune system during parasitism and development. Both this characterisation and the wider understanding of the full *M. hyperodae* venom complement are beneficial in better understanding this biocontrol system. Parasitism dissections on ASW have not detected an increase in encapsulation of *M. hyperodae* eggs (Tomasetto et al., 2017) despite this being one of the most common parasitoid resistance mechanisms documented (Colinet et al., 2013). This may be explained by the multiple *M. hyperodae* venom components working in tandem to prevent such a response, such as in the regulation of autophagy and the PPO cascade, as well as the potential role of calreticulin in the follicle cells of the ovaries.

Parasitised ASW also have their internal organs consumed by the developing egg, though the posterior of the digestive system is left intact (Loan & Lloyd, 1974), and have significantly reduced flight capacity (Goldson et al., 1999). Consumption of host tissues likely explains reduced flight capability, with enzymes such as lipases and proteases allowing for the mobilisation of host nutrients. After parasitism ASW have regressed ovaries and a reduction in testes size (Loan & Lloyd, 1974), leading to reduced fecundity (Barratt et al., 1996). The cause of this reproductive sterilisation is not known, with no currently identified venom components demonstrating a clear link to reproductive manipulation.

The parthenogenetic reproduction of *M. hyperodae* is critical to better understand if the efficacy of biocontrol is to be maintained or improved. There is no indication so far of what is causing *M. hyperodae* to reproduce asexually, though preliminary investigation of the microbiome has not yet revealed parthenogenesis causing endosymbionts, such as Wolbachia. The meiosis-specific genes and their expected expression pattern imply that parthenogenetic reproduction in *M. hyperodae* is either automictic, involving meiosis, or evolved from a mechanism involving meiosis without sufficient time for pseudogenisation or loss of meiosis-specific genes. *M. hyperodae* asexual reproduction may therefore be facultative, which fits with previous reports of impotent male *M. hyperodae* (Goldson et al., 1990). Given previous detection of heterozygous loci in the *M. hyperodae* genome (Iline & Phillips, 2004) it should be assumed that if parthenogenesis is automictic as this data suggests, *M. hyperodae* must rely on a strategy to restore diploidy that maintains at least some heterozygosity.

Such strategies involve either fusion of two haploid meiosis II products (which retains some heterozygosity), the fusion of two meiosis I products (which retains all heterozygosity) or duplication of the entire genome before meiosis (premeiotic duplication, which retains all heterozygosity). The upregulation of genes involved in endoreduplication in the ovaries, which increases cell ploidy, provides a putative mechanism for this premeiotic duplication, though endoreduplication plays an important role in nurse and follicle cells in Drosophila, so expression patterns would need to be investigated to confirm a role in *M. hyperodae* automixis.

## Conclusions

With the first genomic analysis of *M. hyperodae* since the onset of biocontrol decline, we have characterised two traits with significant roles in the initial success (venom) or eventual failure (parthenogenesis) of this biocontrol system. Investigating gene expression across multiple tissues expanded the *M. hyperodae* venom complement to 83 genes while also revealing that calreticulin is not venom specific. Expression localised in the follicle cells of the ovaries may result in the deposition of calreticulin on the egg exochorion to protect the egg in the host. We also show that core meiosis genes are present in the *M. hyperodae* transcriptome and have expression patterns consistent with conserved function. This implies that *M. hyperodae* parthenogenesis is automictic and that they have likely retained the potential for sexual reproduction.

## Acknowledgements

The work was funded by a grant to PKD from Project 2 of the Bioprotection Research Centre and Project 2.2 of Bioprotection Aotearoa (both are Royal Society of New Zealand-funded Centres of Research Excellence). PKD is also funded by Genomics Aotearoa. The authors would like to thank Prof Stephen Goldson, Malvika Bana and Lokesh Kumar for sample collection and P.M. Dearden for critical reading of this manuscript.

## Author Contributions

SNI; Sample collection, molecular biology, bioinformatics, in-situ hybridisation, imaging, manuscript drafting. TWRH; Bioinformatics support and analysis. PKD; Funding, project conception, supervision, manuscript drafting.

## Data availability

All samples used in the analysis (excluding the contaminated Mh_venom_3) are hosted at the National Center for Biotechnology Information Sequence Read Archive (NCBI SRA) database with the accession PRJNA841753. The length-filtered transcriptome assembly is hosted at the NCBI Transcriptome Shotgun Assembly (TSA) database under the same accession. This assembly has a reduced number of transcripts, after those with UniVec sequencing primer hit or those labelled as contaminated during the submission process were removed. Scripts used for analyses are hosted in GitHub repositories as specified in the methods.

## References

Altschul, S. F., Gish, W., Miller, W., Myers, E. W., & Lipman, D. J. (1990). Basic local alignment search tool. J. Mol. Biol., 215(3), 403–410. https://doi.org/10.1016/S0022-2836(05)80360-2

Arakaki, N., Noda, H., & Yamagishi, K. (2000). Wolbachia-induced parthenogenesis in the egg parasitoid Telenomus nawai. Entomol. Exp. Appl., 96(2), 177–184. https://doi.org/10.1046/j.1570-7458.2000.00693.x

Asgari, S., & Rivers, D. B. (2011). Venom proteins from endoparasitoid wasps and their role in host-parasite interactions. Annu. Rev. Entomol., 56, 313–335. https://doi.org/10.1146/annurev-ento-120709-144849

Ashburner, M., Ball, C. A., Blake, J. A., Botstein, D., Butler, H., Cherry, J. M., Davis, A. P., Dolinski, K., Dwight, S. S., Eppig, J. T., Harris, M. A., Hill, D. P., Issel-Tarver, L., Kasarskis, A., Lewis, S., Matese, J. C., Richardson, J. E., Ringwald, M., Rubin, G. M., & Sherlock, G. (2000). Gene ontology: Tool for the unification of biology. Nat. Genet., 25(1), 25–29. https://doi.org/10.1038/75556

Barker, G. M., & Addison, P. J. (2006). Early impact of endoparasitoid Microctonus hyperodae (Hymenoptera: braconidae) after its establishment in Listronotus bonariensis (Coleoptera: Curculionidae) populations of northern New Zealand pastures. J. Econ. Entomol., 99(2), 273–287. https://doi.org/10.1093/jee/99.2.273

Barratt, B. I. P., Evans, A. A., & Johnstone, P. D. (1996). Effect of the ratios of Listronotus bonariensis and Sitona discoideus (Coleoptera: Curculionidae) to their respective parasitoids Microctonus hyperodae and M. aethiopoides (Hymenoptera: Braconidae), on parasitism, host oviposition and feeding in the laboratory. Bull. Entomol. Res., 86(2), 101–108. https://doi.org/10.1017/s0007485300052329

Becchimanzi, A., Avolio, M., Di Lelio, I., Marinelli, A., Varricchio, P., Grimaldi, A., de Eguileor, M., Pennacchio, F., & Caccia, S. (2017). Host regulation by the ectophagous parasitoid wasp Bracon nigricans. J. Insect Physiol., 101, 73–81. https://doi.org/10.1016/j.jinsphys.2017.07.002

Boutet, E., Lieberherr, D., Tognolli, M., Schneider, M., & Bairoch, A. (2007). UniProtKB/Swiss-Prot. Plant Bioinforma., 406, 89–112. https://doi.org/10.1007/978-1-59745-535-0_4

Bryant, D. M., Johnson, K., DiTommaso, T., Tickle, T., Couger, M. B., Payzin-Dogru, D., Lee, T. J., Leigh, N. D., Kuo, T. H., Davis, F. G., Bateman, J., Bryant, S., Guzikowski, A. R., Tsai, S. L., Coyne, S., Ye, W. W., Freeman, R. M., Peshkin, L., Tabin, C. J., … Whited, J. L. (2017). A Tissue-Mapped Axolotl De Novo Transcriptome Enables Identification of Limb Regeneration Factors. Cell Rep., 18(3), 762–776. https://doi.org/10.1016/j.celrep.2016.12.063

Bushnell, B. (2014). BBMap: A Fast, Accurate, Splice-Aware Aligner. https://www.osti.gov/biblio/1241166

Bushnell, B., Rood, J., & Singer, E. (2017). BBMerge – Accurate paired shotgun read merging via overlap. PLoS One, 12(10), e0185056. https://doi.org/10.1371/journal.pone.0185056

Calvete, J. J., Fasoli, E., Sanz, L., Boschetti, E., & Righetti, P. G. (2009). Exploring the venom proteome of the western diamondback rattlesnake, Crotalus atrox, via snake venomics and combinatorial peptide ligand library approaches. J. Proteome Res., 8(6), 3055–3067. https://doi.org/10.1021/pr900249q

Casanovas, P., Goldson, S. L., & Tylianakis, J. M. (2018). Asymmetry in reproduction strategies drives evolution of resistance in biological control systems. PLoS One, 13(12), e0207610. https://doi.org/10.1371/journal.pone.0207610

Choi, H. M. T., Calvert, C. R., Husain, N., Huss, D., Barsi, J. C., Deverman, B. E., Hunter, R. C., Kato, M., Lee, S. M., Abelin, A. C. T., Rosenthal, A. Z., Akbari, O. S., Li, Y., Hay, B. A., Sternberg, P. W., Patterson, P. H., Davidson, E. H., Mazmanian, S. K., Prober, D. A., … Pierce, N. A. (2016). Mapping a multiplexed zoo of mRNA expression. Dev., 143(19), 3632–3637. https://doi.org/10.1242/dev.140137

Choi, H. M. T., Schwarzkopf, M., Fornace, M. E., Acharya, A., Artavanis, G., Stegmaier, J., Cunha, A., & Pierce, N. A. (2018). Third-generation in situ hybridization chain reaction: Multiplexed, quantitative, sensitive, versatile, robust. Dev., 145(12), dev165753. https://doi.org/10.1242/dev.165753

Colinet, D., Cazes, D., Belghazi, M., Gatti, J. L., & Poirié, M. (2011). Extracellular superoxide dismutase in insects: Characterization, function, and interspecific variation in parasitoid wasp venom. J. Biol. Chem., 286(46), 40110–40121. https://doi.org/10.1074/jbc.M111.288845

Colinet, D., Deleury, E., Anselme, C., Cazes, D., Poulain, J., Azéma-Dossat, C., Belghazi, M., Gatti, J. L., & Poirié, M. (2013). Extensive inter- and intraspecific venom variation in closely related parasites targeting the same host: The case of Leptopilina parasitoids of Drosophila. Insect Biochem. Mol. Biol., 43(7), 601–611. https://doi.org/10.1016/j.ibmb.2013.03.010

Cônsoli, F. L., Brandt, S. L., Coudron, T. A., & Vinson, S. B. (2005). Host regulation and release of parasitism-specific proteins in the system Toxoneuron nigriceps-Heliothis virescens. Comp. Biochem. Physiol. - B Biochem. Mol. Biol., 142(2), 181–191. https://doi.org/10.1016/j.cbpc.2005.07.002

Crawford, A. M., Brauning, R., Smolenski, G., Ferguson, C. M., Barton, D. M., Wheeler, T. T., & Mcculloch, A. (2008). The constituents of Microctonus sp. parasitoid venoms. Insect Mol. Biol., 17(3), 313–324. https://doi.org/10.1111/j.1365-2583.2008.00802.x

Crawford, A. M., Still, L. A., & Smith, P. (2006). A histological examination of the venom apparatus of Microctonus hyperodae loan (Hymenoptera: Braconidae). New Zeal. Entomol., 29(1), 103–105. https://doi.org/10.1080/00779962.2006.9722144

Dalmon, A., Gayral, P., Decante, D., Klopp, C., Bigot, D., Thomasson, M., Aherniou, E., Alaux, C., & Conte, Y. Le. (2019). Viruses in the invasive hornet Vespa velutina. Viruses, 11(11), 1041. https://doi.org/10.3390/v11111041

Danneels, E. L., Rivers, D. B., & de Graaf, D. C. (2010). Venom proteins of the parasitoid wasp Nasonia vitripennis: Recent discovery of an untapped pharmacopeia. In Toxins (Vol. 2, Issue 4, pp. 494–516). https://doi.org/10.3390/toxins2040494

Dorémus, T., Jouan, V., Urbach, S., Cousserans, F., Wincker, P., Ravallec, M., Wajnberg, E., & Volkoff, A. N. (2013). Hyposoter didymator uses a combination of passive and active strategies to escape from the Spodoptera frugiperda cellular immune response. J. Insect Physiol., 59(4), 500–508. https://doi.org/10.1016/j.jinsphys.2013.02.010

Drezen, J. M., Leobold, M., Bézier, A., Huguet, E., Volkoff, A. N., & Herniou, E. A. (2017). Endogenous viruses of parasitic wasps: variations on a common theme. Curr. Opin. Virol., 25, 41–48. https://doi.org/10.1016/j.coviro.2017.07.002

Finn, R. D., Bateman, A., Clements, J., Coggill, P., Eberhardt, R. Y., Eddy, S. R., Heger, A., Hetherington, K., Holm, L., Mistry, J., Sonnhammer, E. L. L., Tate, J., & Punta, M. (2014). Pfam: The protein families database. In Nucleic Acids Research. https://doi.org/10.1109/TCSVT.2017.2671899

Finn, R. D., Clements, J., & Eddy, S. R. (2011). HMMER web server: Interactive sequence similarity searching. Nucleic Acids Res., 39(SUPPL. 2). https://doi.org/10.1093/nar/gkr367

Gettins, P. G. W. (2002). Serpin structure, mechanism, and function. Chem. Rev., 102(12), 4751–4804.

Goldson, S. L., McNeill, M. R., Proffitt, J. R., Barker, G. M., Addison, P. J., Barratt, B. I. P., & Ferguson, C. M. (1993). Systematic mass rearing and release of Microctonus hyperodae (Hym.: Braconidae, Euphorinae), a parasitoid of the argentine stem weevil Listronotus bonariensis (Col.: Curculionidae) and records of its establishment in New Zealand. Entomophaga, 38(4), 527–536. https://doi.org/10.1007/BF02373087

Goldson, S. L., McNeill, M. R., Stufkens, M. W., Proffitt, J. R., Pottinger, R. P., & Farrell, J. A. (1990). Importation and quarantine of Microctonus hyperodae, a South American parasitoid of Argentine stem weevil. In Proceedings of the Forty Third New Zealand Weed and Pest Control Conference (pp. 334–338). New Zealand Weed and Pest Control Society Inc.

Goldson, S. L., Proffitt, J. R., & Baird, D. B. (1999). Listronotus bonariensis (Coleoptera: Curculionidae) flight in Canterbury, New Zealand. Bull. Entomol. Res., 89(5), 423–431. https://doi.org/10.1017/s0007485399000553

Goldson, S. L., & Tomasetto, F. (2016). Apparent acquired resistance by a weevil to its parasitoid is influenced by host plant. Front. Plant Sci., 7(AUG2016). https://doi.org/10.3389/fpls.2016.01259

Grabherr, M. G., Haas, B. J., Yassour, M., Levin, J. Z., Thompson, D. A., Amit, I., Adiconis, X., Fan, L., Raychowdhury, R., Zeng, Q., Chen, Z., Mauceli, E., Hacohen, N., Gnirke, A., Rhind, N., Di Palma, F., Birren, B. W., Nusbaum, C., Lindblad-Toh, K., … Regev, A. (2011). Full-length transcriptome assembly from RNA-Seq data without a reference genome. Nat. Biotechnol., 29(7), 644–652. https://doi.org/10.1038/nbt.1883

Hagiwara, K., Kikuchi, T., Endo, Y., Huqun, Usui K., Takahashi, M., Shibata, N., Kusakabe, T., Xin, H., Hoshi, S., Miki, M., Inooka, N., Tokue, Y., & Nukiwa, T. (2003). Mouse SWAM1 and SWAM2 Are Antibacterial Proteins Composed of a Single Whey Acidic Protein Motif. J. Immunol., 170(4), 1973–1979. https://doi.org/10.4049/jimmunol.170.4.1973

Harrop, T. W. R., Le Lec, M. F., Jauregui, R., Taylor, S. E., Inwood, S. N., van Stijn, T., Henry, H., Skelly, J. G., Ganesh, S., Ashby, R. L., Jacobs, J. M. E., Goldson, S. L., & Dearden, P. K. (2020). Genetic diversity in invasive populations of argentine stem weevil associated with adaptation to biocontrol. Insects, 11(7), 1–14. https://doi.org/10.3390/insects11070441

Heavner, M. E., Gueguen, G., Rajwani, R., Pagan, P. E., Small, C., & Govind, S. (2013). Partial venom gland transcriptome of a Drosophila parasitoid wasp, Leptopilina heterotoma, reveals novel and shared bioactive profiles with stinging Hymenoptera. Gene, 526(2), 195–204. https://doi.org/10.1016/j.gene.2013.04.080

Highfield, A., Kevill, J., Kevill, J., Mordecai, G., Mordecai, G., Hunt, J., Henderson, S., Sauvard, D., Feltwell, J., Martin, S. J., Sumner, S., Schroeder, D. C., Schroeder, D. C., & Schroeder, D. C. (2020). Detection and replication of moku virus in honey bees and social wasps. Viruses, 12(6), 607. https://doi.org/10.3390/v12060607

Iline, I. I., & Phillips, C. B. (2004). Allozyme markers to help define the South American origins of Microctonus hyperodae (Hymenoptera: Braconidae) established in New Zealand for biological control of Argentine stem weevil. Bull. Entomol. Res., 94(3), 229–234. https://doi.org/10.1079/ber2004303

Kanehisa, M., Goto, S., Sato, Y., Furumichi, M., & Tanabe, M. (2012). KEGG for integration and interpretation of large-scale molecular data sets. Nucleic Acids Res., 40(D1). https://doi.org/10.1093/nar/gkr988

Kanost, M. R., & Gorman, M. J. (2008). Phenoloxidases in Insect Immunity. Insect Immunol., 1, 69–96. https://doi.org/10.1016/B978-012373976-6.50006-9

Kolde, R. (2019). Pretty Heatmaps (R package version 1.0.12; pp. 1–8). https://cran.r-project.org/package=pheatmap

Kongton, K., McCall, K., & Phongdara, A. (2014). Identification of gamma-interferon-inducible lysosomal thiol reductase (GILT) homologues in the fruit fly Drosophila melanogaster. Dev. Comp. Immunol., 44(2), 389–396. https://doi.org/10.1016/j.dci.2014.01.007

Köster, J., & Rahmann, S. (2012). Snakemake-a scalable bioinformatics workflow engine. Bioinformatics, 28(19), 2520–2522. https://doi.org/10.1093/bioinformatics/bts480

Koziy, R. V., Wood, S. C., Kozii, I. V., van Rensburg, C. J., Moshynskyy, I., Dvylyuk, I., & Simko, E. (2019). Deformed Wing Virus Infection in Honey Bees (Apis mellifera L.). Vet. Pathol., 56(4), 636–641. https://doi.org/10.1177/0300985819834617

Kramer, K. J., & Koga, D. (1986). Insect chitin. Physical state, synthesis, degradation and metabolic regulation. Insect Biochem., 16(6), 851–877. https://doi.org/10.1016/0020-1790(86)90059-4

Laurino, S., Grossi, G., Pucci, P., Flagiello, A., Bufo, S. A., Bianco, G., Salvia, R., Vinson, S. B., Vogel, H., & Falabella, P. (2016). Identification of major Toxoneuron nigriceps venom proteins using an integrated transcriptomic/proteomic approach. Insect Biochem. Mol. Biol., 76, 49–61. https://doi.org/10.1016/j.ibmb.2016.07.001

Lilly, M. A., & Sptadling, A. C. (1996). The Drosophila endocycle is controlled by cyclin E and lacks a checkpoint ensuring S-phase completion. Genes Dev., 10(19), 2514–2526. https://doi.org/10.1101/gad.10.19.2514

Lin, H., & Spradling, A. C. (1997). A novel group of pumilio mutations affects the asymmetric division of germline stem cells in the Drosophila ovary. Development, 124(12), 2463–2476. https://doi.org/10.1242/dev.124.12.2463

Lin, Z., Wang, R. J., Cheng, Y., Du, J., Volovych, O., Han, L. Bins, Li, J. C., Hu, Y., Lu, Z. Y., Lu, Z., & Zou, Z. (2019). Insights into the venom protein components of Microplitis mediator, an endoparasitoid wasp. Insect Biochem. Mol. Biol., 105, 33–42. https://doi.org/10.1016/j.ibmb.2018.12.013

Liu, N. Y., Huang, J. M., Ren, X. M., Xu, Z. W., Yan, N. S., & Zhu, J. Y. (2018). Superoxide dismutase from venom of the ectoparasitoid Scleroderma guani inhibits melanization of hemolymph. Arch. Insect Biochem. Physiol., 99(3), e21503. https://doi.org/10.1002/arch.21503

Liu, N. Y., Xu, Z. W., Yan, W., Ren, X. M., Zhang, Z. Q., & Zhu, J. Y. (2018). Venomics reveals novel ion transport peptide-likes (ITPLs) from the parasitoid wasp Tetrastichus brontispae. Toxicon, 141, 88–93. https://doi.org/10.1016/j.toxicon.2017.11.008

Loan, C. C., & Lloyd, D. C. (1974). Description and field biology of Microctonus hyperodae Loan, n. sp. [Hymenoptera: Braconidae, Euphorinae] a parasite of Hyperodes bonariensis in South America [Coleoptera: Curculionidae]. Entomophaga, 19(1), 7–12. https://doi.org/10.1007/BF02371504

Love, M. I., Huber, W., & Anders, S. (2014). Moderated estimation of fold change and dispersion for RNA-seq data with DESeq2. Genome Biol., 15(12), 550. https://doi.org/10.1186/s13059-014-0550-8

Ma, W. J., & Schwander, T. (2017). Patterns and mechanisms in instances of endosymbiont-induced parthenogenesis. J. Evol. Biol., 30(5), 868–888. https://doi.org/10.1111/jeb.13069

Maines, J. Z., Stevens, L. M., Tong, X., & Stein, D. (2004). Drosophila dMyc is required for ovary cell growth and endoreplication. Development. 2004;131(4):775–86. doi: 10.1242/dev.00932

Mordecai, G. J., Brettell, L. E., Pachori, P., Villalobos, E. M., Martin, S. J., Jones, I. M., & Schroeder, D. C. (2016). Moku virus; A new Iflavirus found in wasps, honey bees and Varroa. Sci. Rep., 6(1), 1–6. https://doi.org/10.1038/srep34983

Moreau, S. J. M., & Asgari, S. (2015). Venom proteins from parasitoid wasps and their biological functions. Toxins (Basel)., 7(7), 2385–2412. https://doi.org/10.3390/toxins7072385

Nair, D. G., Fry, B. G., Alewood, P., Kumar, P. P., & Kini, R. M. (2007). Antimicrobial activity of omwaprin, a new member of the waprin family of snake venom proteins. Biochem. J., 402(1), 93–104. https://doi.org/10.1042/BJ20060318

Nakamatsu, Y., & Tanaka, T. (2004). Venom of Euplectrus separatae causes hyperlipidemia by lysis of host fat body cells. J. Insect Physiol., 50(4), 267–275. https://doi.org/10.1016/j.jinsphys.2003.12.005

Parkinson, N., Conyers, C., & Smith, I. (2002). A venom protein from the endoparasitoid wasp Pimpla hypochondriaca is similar to snake venom reprolysin-type metalloproteases. J. Invertebr. Pathol., 79(2), 129–131. https://doi.org/10.1016/S0022-2011(02)00033-2

Patro, R., Duggal, G., Love, M. I., Irizarry, R. A., & Kingsford, C. (2017). Salmon provides fast and bias-aware quantification of transcript expression. Nat. Methods, 14(4), 417–419. https://doi.org/10.1038/nmeth.4197

Perkin, L. C., Friesen, K. S., Flinn, P. W., & Oppert, B. (2015). Venom gland components of the ectoparasitoid wasp, Anisopteromalus calandrae. J. Venom Res., 6, 19–37. http://www.ncbi.nlm.nih.gov/pubmed/26998218 %0A http://www.pubmedcentral.nih.gov/articlerender.fcgi?artid=PMC4776022

Petersen, T. N., Brunak, S., Von Heijne, G., & Nielsen, H. (2011). SignalP 4.0: Discriminating signal peptides from transmembrane regions. Nat. Methods, 8(10), 785–786. https://doi.org/10.1038/nmeth.1701

Phillips, C. B. (1995). Intraspecific variation in Microctonus hyperodae and M. aethiopoides (Hymenoptera: Braconidae); significance for their use of biological control agents. Lincoln University.

Popay, A. J., McNeill, M. R., Goldson, S. L., & Ferguson, C. M. (2011). The current status of Argentine stem weevil (Listronotus bonariensis) as a pest in the North Island of New Zealand. New Zeal. Plant Prot., 64, 55–62. https://doi.org/10.30843/nzpp.2011.64.5962

Powell, S., Szklarczyk, D., Trachana, K., Roth, A., Kuhn, M., Muller, J., Arnold, R., Rattei, T., Letunic, I., Doerks, T., Jensen, L. J., Von Mering, C., & Bork, P. (2012). eggNOG v3.0: Orthologous groups covering 1133 organisms at 41 different taxonomic ranges. Nucleic Acids Res., 40(D1), D284–D289. https://doi.org/10.1093/nar/gkr1060

Prestidge, R. A., Barker, G. M., & Pottinger, R. P. (1991). The economic cost of Argentine stem weevil in pastures in New Zealand. In Proceedings of the New Zealand Weed and Pest Control Conference (Vol. 44, pp. 165–170). New Zealand Weed and Pest Control Society Inc. https://doi.org/10.30843/nzpp.1991.44.10825

Price, D. R. G., Bell, H. A., Hinchliffe, G., Fitches, E., Weaver, R., & Gatehouse, J. A. (2009). A venom metalloproteinase from the parasitic wasp Eulophus pennicornis is toxic towards its host, tomato moth (Lacanobia oleracae). Insect Mol. Biol., 18(2), 195–202. https://doi.org/10.1111/j.1365-2583.2009.00864.x

Pruitt, K. D., Tatusova, T., & Maglott, D. R. (2007). NCBI reference sequences (RefSeq): A curated non-redundant sequence database of genomes, transcripts and proteins. Nucleic Acids Res., 35(SUPPL. 1), D501–D504. https://doi.org/10.1093/nar/gkl842

R Development Core Team. (2020). A Language and Environment for Statistical Computing. R Found. Stat. Comput., https://www.R-project.org. http://www.r-project.org

Ramesh, M. A., Malik, S. B., & Logsdon, J. M. (2005). A phylogenomic inventory of meiotic genes: Evidence for sex in Giardia and an early eukaryotic origin of meiosis. Curr. Biol., 15(2), 185–191. https://doi.org/10.1016/j.cub.2005.01.003

Rivers, D. B., & Denlinger, D. L. (1995). Venom-induced alterations in fly lipid metabolism and its impact on larval development of the ectoparasitoid Nasonia vitripennis (Walker)(Hymenoptera: Pteromalidae). J. Invertebr. Pathol., 66(2), 104–110.

Sahu, A., & Birner-Gruenberger, R. (2013). Lipases. Encycl. Met. https://doi.org/10.1007/978-1-4614-1533-6

Saikhedkar, N., Summanwar, A., Joshi, R., & Giri, A. (2015). Cathepsins of lepidopteran insects: Aspects and prospects. Insect Biochem. Mol. Biol., 64, 51–59. https://doi.org/https://doi.org/10.1016/j.ibmb.2015.07.005

Schurko, A. M., & Logsdon, J. M. (2008). Using a meiosis detection toolkit to investigate ancient asexual “scandals” and the evolution of sex. BioEssays, 30(6), 579–589. https://doi.org/10.1002/bies.20764

Schurko, A. M., Mazur, D. J., & Logsdon Jr, J. M. (2010). Inventory and phylogenomic distribution of meiotic genes in Nasonia vitripennis and among diverse arthropods. Insect Mol. Biol., 19, 165–180.

Scieuzo, C., Salvia, R., Franco, A., Pezzi, M., Cozzolino, F., Chicca, M., Scapoli, C., Vogel, H., Monti, M., Ferracini, C., Pucci, P., Alma, A., & Falabella, P. (2021). An integrated transcriptomic and proteomic approach to identify the main Torymus sinensis venom components. Sci. Rep., 11(1), 1–25. https://doi.org/10.1038/s41598-021-84385-5

Shen, W., & Ren, H. (2021). TaxonKit: A practical and efficient NCBI taxonomy toolkit. J. Genet. Genomics. https://doi.org/10.1016/j.jgg.2021.03.006

Siebert, A. L., Wheeler, D., & Werren, J. H. (2015). A new approach for investigating venom function applied to venom calreticulin in a parasitoid wasp. Toxicon, 107, 304–316. https://doi.org/10.1016/j.toxicon.2015.08.012

Sim, A. D., & Wheeler, D. (2016). The venom gland transcriptome of the parasitoid wasp Nasonia vitripennis highlights the importance of novel genes in venom function. BMC Genomics, 17(1), 1–16. https://doi.org/10.1186/s12864-016-2924-7

Simão, F. A., Waterhouse, R. M., Ioannidis, P., Kriventseva, E. V., & Zdobnov, E. M. (2015). BUSCO: Assessing genome assembly and annotation completeness with single-copy orthologs. Bioinformatics, 31(19), 3210–3212. https://doi.org/10.1093/bioinformatics/btv351

Soneson, C., Love, M. I., & Robinson, M. D. (2016). Differential analyses for RNA-seq: Transcript-level estimates improve gene-level inferences. F1000Research, 4. https://doi.org/10.12688/F1000RESEARCH.7563.2

Stouthamer, R., Luck, R. F., & Hamilton, W. D. (1990). Antibiotics cause parthenogenetic Trichogramma (Hymenoptera/trichogrammatidae) to revert to sex. Proc. Natl. Acad. Sci. U. S. A., 87(7), 2424–2427. https://doi.org/10.1073/pnas.87.7.2424

Teng, Z., Wu, H., Ye, X., Fang, Q., Zhou, H., & Ye, G. (2021). An Endoparasitoid Uses Its Egg Surface Proteins to Regulate Its Host Immune Response. Insect Sci. https://doi.org/10.1111/1744-7917.12978

Thomas, P., & Asgari, S. (2011). Inhibition of melanization by a parasitoid serine protease homolog venom protein requires both the clip and the non-catalytic protease-like domains. Insects, 2(4), 509–514. https://doi.org/10.3390/insects2040509

Tomasetto, F., Tylianakis, J. M., Reale, M., Wratten, S. D., & Goldson, S. L. (2017). Intensified agriculture favors evolved resistance to biological control. Proc. Natl. Acad. Sci. U. S. A., 114(15), 3885–3890. https://doi.org/10.1073/pnas.1618416114

Tvedte, E. S., Forbes, A. A., Logsdon, J. M., & Reed, F. (2017). Retention of Core Meiotic Genes Across Diverse Hymenoptera. J. Hered., 108(7), 791–806. https://doi.org/10.1093/jhered/esx062

Tvedte, E. S., Logsdon, J. M., & Forbes, A. A. (2019). Sex loss in insects: causes of asexuality and consequences for genomes. Curr. Opin. Insect Sci., 31, 77–83. https://doi.org/10.1016/j.cois.2018.11.007

Vaiyapuri, S., Wagstaff, S. C., Watson, K. A., Harrison, R. A., Gibbins, J. M., & Hutchinson, E. G. (2010). Purification and functional characterisation of rhiminopeptidase A, a novel aminopeptidase from the venom of Bitis gabonica rhinoceros. PLoS Negl. Trop. Dis., 4(8), e796. https://doi.org/10.1371/journal.pntd.0000796

Wang, L., Fang, Q., Qian, C., Wang, F., Yu, X. Q., & Ye, G. (2013). Inhibition of host cell encapsulation through inhibiting immune gene expression by the parasitic wasp venom calreticulin. Insect Biochem. Mol. Biol., 43(10), 936–946. https://doi.org/10.1016/j.ibmb.2013.07.010

Webster, S. G., Keller, R., & Dircksen, H. (2012). The CHH-superfamily of multifunctional peptide hormones controlling crustacean metabolism, osmoregulation, moulting, and reproduction. In General and Comparative Endocrinology (Vol. 175, Issue 2, pp. 217–233). https://doi.org/10.1016/j.ygcen.2011.11.035

Wilfert, L., Long, G., Leggett, H. C., Schmid-Hempel, P., Butlin, R., Martin, S. J. M., & Boots, M. (2016). Honeybee disease: Deformed wing virus is a recent global epidemic in honeybees driven by Varroa mites. Science (80-.)., 351(6273), 594–597. https://doi.org/10.1126/science.aac9976

Wood, D. E., Lu, J., & Langmead, B. (2019). Improved metagenomic analysis with Kraken 2. Genome Biol., 20(1), 1–13. https://doi.org/10.1186/s13059-019-1891-0

Yan, Z., Fang, Q., Liu, Y., Xiao, S., Yang, L., Wang, F., An, C., Werren, J. H., & Ye, G. (2017). A venom serpin splicing isoform of the endoparasitoid wasp Pteromalus puparum suppresses host prophenoloxidase cascade by forming complexes with host hemolymph proteinases. J. Biol. Chem., 292(3), 1038–1051. https://doi.org/10.1074/jbc.M116.739565

Yan, Z., Fang, Q., Wang, L., Liu, J., Zhu, Y., Wang, F., Li, F., Werren, J. H., & Ye, G. (2016). Insights into the venom composition and evolution of an endoparasitoid wasp by combining proteomic and transcriptomic analyses. Sci. Rep., 6. https://doi.org/10.1038/srep19604

Yang, L., Yang, Y., Liu, M. M., Yan, Z. C., Qiu, L. M., Fang, Q., Wang, F., Werren, J. H., & Ye, G. Y. (2020). Identification and Comparative Analysis of Venom Proteins in a Pupal Ectoparasitoid, Pachycrepoideus vindemmiae. Front. Physiol., 11, 9. https://doi.org/10.3389/fphys.2020.00009

Ye, X. Qian, Shi, M., Huang, J. Hua, & Chen, X. Xin. (2018). Parasitoid polydnaviruses and immune interaction with secondary hosts. Dev. Comp. Immunol., 83, 124–129. https://doi.org/10.1016/j.dci.2018.01.007

Yu, G., Wang, L. G., Han, Y., & He, Q. Y. (2012). ClusterProfiler: An R package for comparing biological themes among gene clusters. Omi. A J. Integr. Biol., 16(5), 284–287. https://doi.org/10.1089/omi.2011.0118

Zhang, G., Lu, Z. Q., Jiang, H., & Asgari, S. (2004). Negative regulation of prophenoloxidase (proPO) activation by a clip-domain serine proteinase homolog (SPH) from endoparasitoid venom. Insect Biochem. Mol. Biol., 34(5), 477–483. https://doi.org/10.1016/j.ibmb.2004.02.009

Zhang, G., Schmidt, O., & Asgari, S. (2006). A calreticulin-like protein from endoparasitoid venom fluid is involved in host hemocyte inactivation. Dev. Comp. Immunol., 30(9), 756–764.

