## Supplemental Figures for "The parthenogenesis mechanism and venom complement of the parasitoid wasp *Microctonus hyperodae*, a declining biocontrol agent"

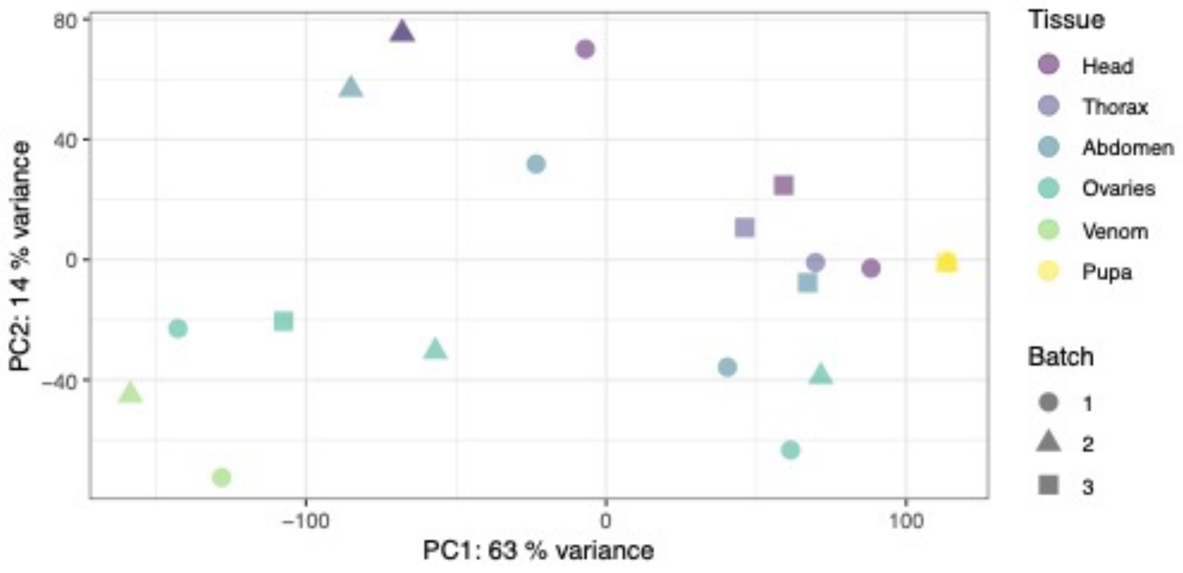


*Supplementary Figure 1 A principal component analysis for M. hyperodae RNA-seq samples. Points are coloured based on tissue identity, and shaped based on the dissection/extraction batch.*

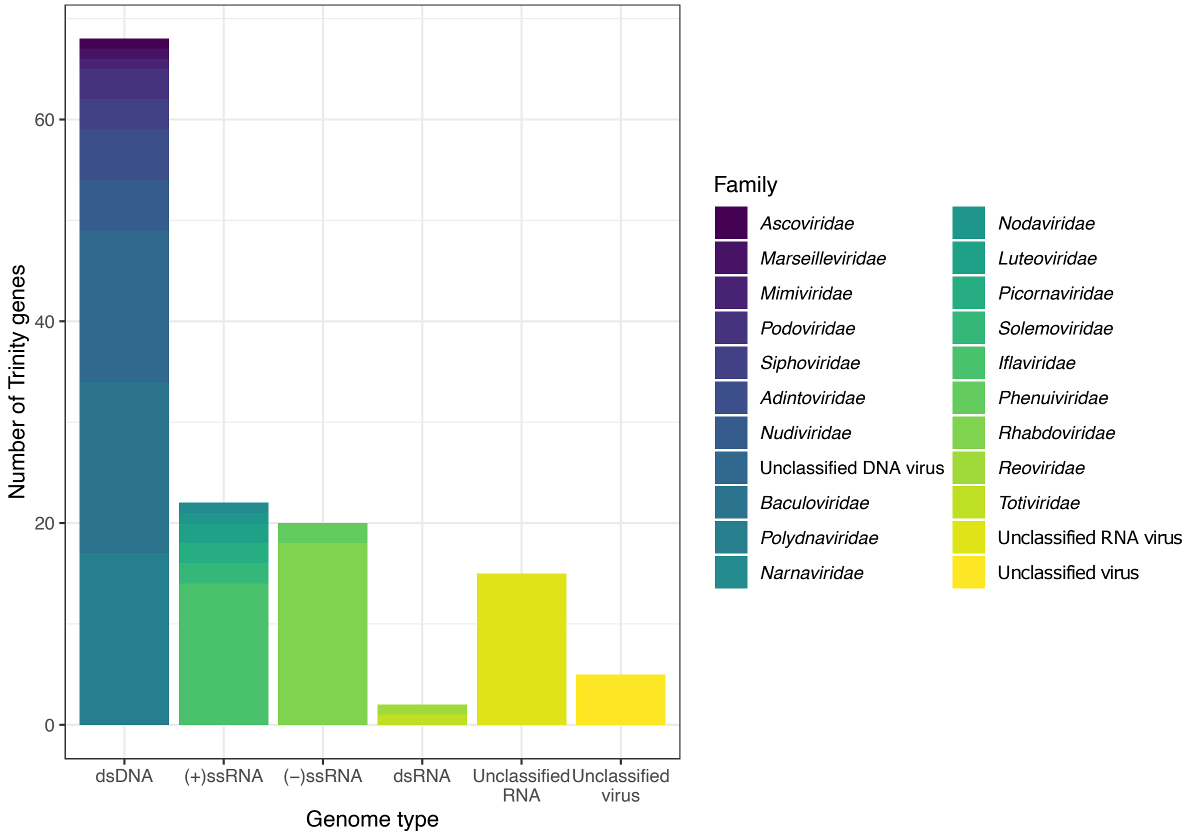


*Supplementary Figure 2 A bar graph showing the number of Trinity genes in the M. hyperodae transcriptome with significant viral BlastX hits from the reciprocal search. Hits are coloured based on viral family, and bars separated by genome type.*

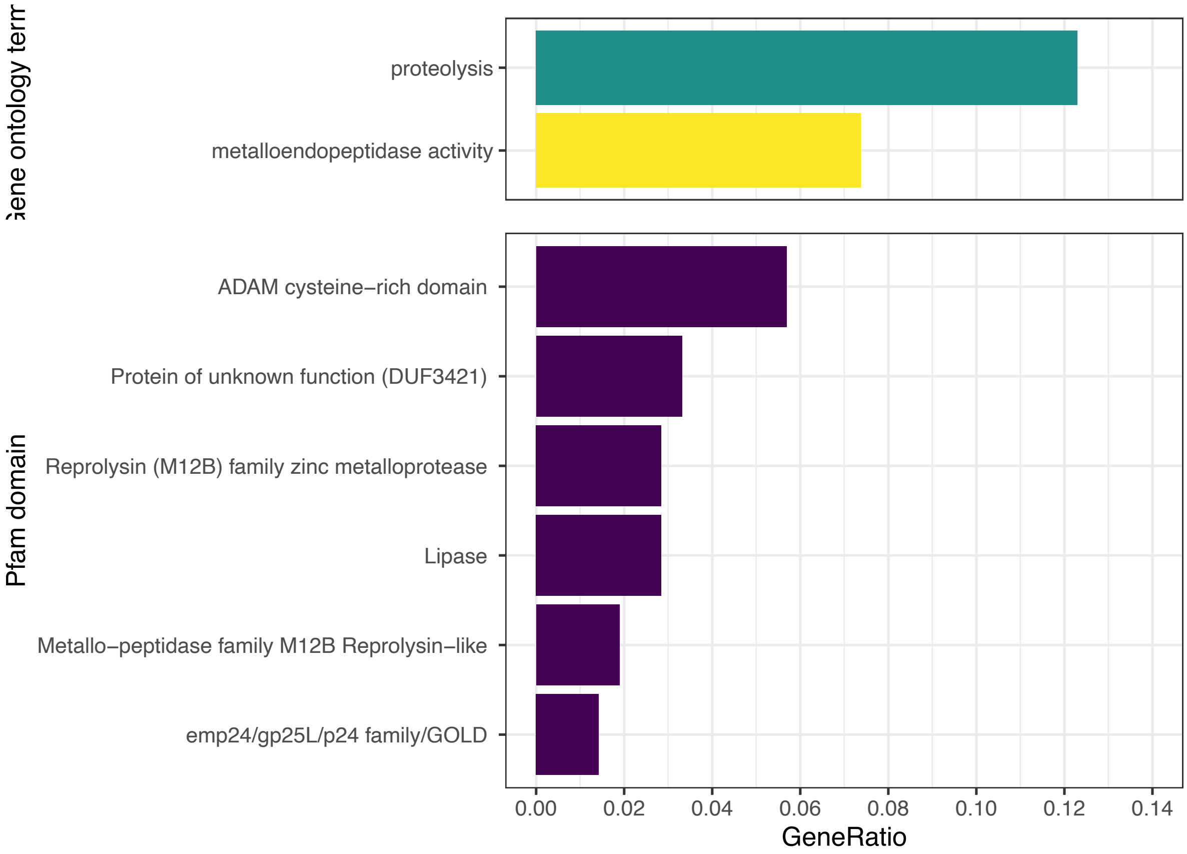


*Supplementary Figure 3 A bar graph displaying significantly enriched gene ontology terms and Pfam protein domains for the M. hyperodae venom differential gene expression, as determined with clusterProfiler. The GeneRatio is the ratio of DEGs with the associated Pfam domain. Significantly enriched terms were identified as those with an adjusted P-value less than 0.05. GO terms are coloured based on GO domain, with biological processes in green and molecular functions in yellow.*

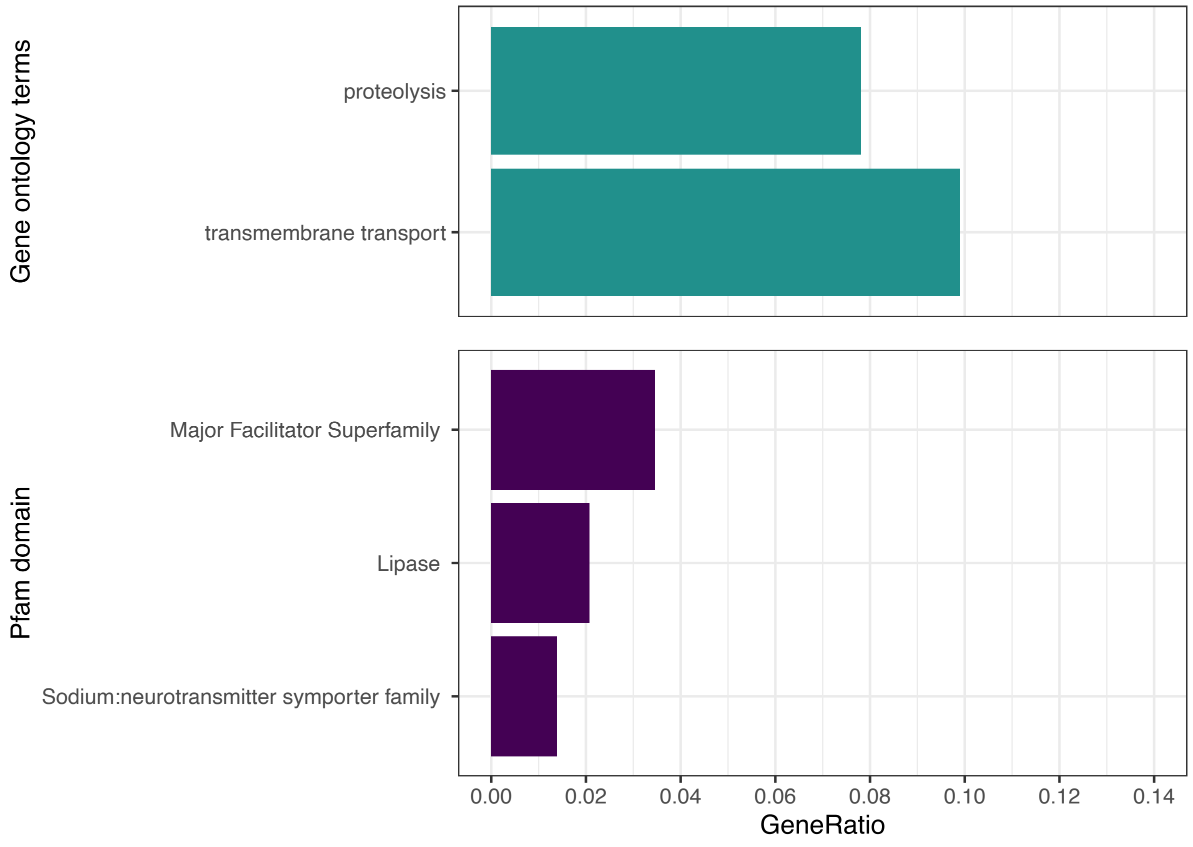


*Supplementary Figure 4 A bar graph displaying significantly enriched gene ontology terms and Pfam protein domains for the M. hyperodae ovary differential gene expression, as determined with clusterProfiler. The GeneRatio is the ratio of DEGs with the associated Pfam domain. Significantly enriched terms were identified as those with an adjusted P-value less than 0.05. GO terms are coloured based on GO domain, with biological processes in green and molecular functions in yellow.*
